# Human vagus nerve fascicular anatomy: a microCT segmentation and histological study

**DOI:** 10.1101/2023.07.04.547643

**Authors:** Nicole Thompson, Svetlana Mastitskaya, Francesco Iacoviello, Paul R. Shearing, Kirill Aristovich, David Holder

## Abstract

**Background:** Previous research has revealed the logical mapping of fascicles in both human somatic and pig vagus nerves, but the organization of fascicles within the human vagus nerve remains largely unknown. Understanding its fascicular arrangement would significantly advance our knowledge of the autonomic nervous system and facilitate studies and application of selective vagus nerve stimulation to avoid off-target effects. The purpose of this study was to trace the thoracic branches of human vagus nerves, investigate their fascicular organization, and analyze the nerves histologically and morphologically.

**Methods:** Both left and right vagus nerves were dissected from human cadavers, preserving the cardiac, recurrent laryngeal, and pulmonary branches. The nerves were prepared, scanned using microCT, and the fascicles segmented and traced from their branching points. Histology and immunohistochemistry were performed for morphological analysis and validation of the microCT segmentation. The data was then analyzed and compared between nerves.

**Results:** The organization of the cardiac, pulmonary, and recurrent laryngeal fascicles was observed for a short distance from their entry point into the nerves. Initially, left vagus nerves showed merging of cardiac and pulmonary fascicles, while the recurrent laryngeal fascicles remained separate. In right vagus nerves, the cardiac fascicles merged with both pulmonary and recurrent laryngeal fascicles. MicroCT imaging limitations prevented visualization and tracing of fiber organization within merged fascicles. Immunohistochemistry and morphological analysis revealed that right vagus nerves were larger and had more fascicles than the left and fascicle counts varied along the nerve, indicating anastomoses. The superior cardiac branch was separate from other fascicles near the VNS cuff placement.

**Conclusions:** It is possible that organ-specific fibers may still retain some spatial organization despite most fascicles being merged at cervical level. However, fiber tracing and *in vivo* studies could provide valuable information beyond microCT to resolve this further. The separate superior cardiac fascicles offer potential for targeted neuromodulation of the heart, benefiting conditions like myocardial infarction, heart failure, and atrial fibrillation. Overall, the study provides insights into the morphology and anatomy of human vagus nerves. Our findings thereby contribute to the development of selective vagus nerve stimulation strategies for more precise autonomic regulation.

## Introduction

The functional anatomy of somatic peripheral nerves has been extensively studied using histological tracing, which has revealed a logical mapping of fascicles to dermatomes and muscle groups (1,2). However, the anatomical relationship and functional organization of fascicles within the human vagus nerve, the main peripheral nerve of the autonomic nervous system (ANS) that innervates numerous visceral organs in the thorax and abdomen, as well as the larynx (3), remains largely unknown (4–6). Understanding the organization of fascicles in the vagus nerve would represent a significant advancement in our knowledge of the functional anatomy of the ANS, providing insights into neurobiological principles and facilitating studies on neural control, neurophysiology, and clinical applications such as nerve repair and vagus nerve stimulation (VNS) (3,7–15).

In particular, the ability to selectively modulate specific target organs or functions using spatially selective vagus nerve cuffs has opened new possibilities for therapeutic interventions. However, achieving such selectivity requires a comprehensive understanding of the fascicular organization within the vagus nerve, specifically at the level of cuff placement, to avoid off-target effects commonly observed, such as cough, dyspnea, and bradycardia (16,17). Currently, there is limited knowledge regarding the fascicular anatomy of the vagus nerve in humans, hindering the development and optimization of spatially selective neuromodulation techniques.

Previous studies have demonstrated reproducible fascicular organization of the left cervical vagus nerves in pigs, in relation to cardiac, pulmonary, and recurrent laryngeal functions. This suggests that fascicles in the ANS are organized, to a significant extent, according to function or organ controlled. If a similar organization is found in humans, it could lead to a paradigm shift in vagus nerve stimulation, allowing selective stimulation and improving therapeutic efficacy by avoiding off-target effects (18–21).

However, there are notable anatomical differences between pig and human vagus nerves, including variations in fascicle count and diameter. While pig vagus nerves contain between 20 and 50 fascicles at the cervical level, humans have an average of 4 to 10 fascicles (22,23),. Additionally, the diameters of fascicles in pig vagus nerves are smaller compared to humans. To date, only a limited number of histological studies have examined cross-sections of human vagus nerves at the cervical level, with no data available on the rest of the nerve or comprehensive tracing of fascicles along its length (1–4).

In this study, the purpose was to trace the thoracic branches, including the cardiac, pulmonary, and recurrent laryngeal fascicles, of human vagus nerves using microCT (n=5, 3 left, 2 right vagi) and investigate the fascicular organization along the length of the nerves. Additionally, we performed histology and immunohistochemistry (IHC) at regular intervals along the vagal trunk and from each nerve branch to gain morphological insights and contribute to our understanding of the human vagus nerve (n=10, 5 left, 5 right). Specifically, we aimed to address the following questions:

1. How are the fascicles of the human vagus nerve arranged, and to what extent do they exhibit organotopic organization, similar to the pig model?
2. What valuable information can be obtained from histological and immunohistochemical analysis of the human vagus nerve?

By answering these questions, we aim to provide comprehensive insights into the fascicular anatomy and organization of the human vagus nerve, contribute to the existing database of human vagus nerves, and enhance our understanding of general nerve morphology, fascicular arrangement and merging patterns within the nerve. Furthermore, this research has implications for improving the clinical applications of VNS, nerve repair, and regeneration, ultimately leading to more precise and effective therapeutic interventions. Additionally, the preliminary investigation of histological and IHC analyses were expected to contribute to further understanding of anatomy and morphology of the vagus nerve and serve as a foundation and step towards future advanced fiber tracing studies.

## Methods

### Human vagus nerve samples

The Evelyn Cambridge Surgical Training Centre, A Cambridge University Health Partners Facility, loaned human vagus nerve samples for this study. The Human Tissue Authority (HTA) Designated Individual (DI), namely Dr C R Constant, approved the loan of human tissue from first-person, signed and witnessed consented (willed) donors for the use in this project. Dissection of vagus nerves from the donors was supervised by the HTA DI for the Centre’s license and all subsequent transport and work on the nerves was pre-approved by the HTA DI on their license and specified in a Material Transfer/Loan Agreement. Separate ethical approval was not required.

### Nerve samples and post-mortem dissection

Both left and right vagus nerves were dissected from frozen human cadavers (N=5, n=10). The nerves were dissected from the cervical region down to the branching regions of cardiac, recurrent laryngeal and pulmonary branches, with all the branches left attached to the main trunk of the vagus nerve. Specifically, vagus nerves were exposed using techniques that are similar to surgery for vagus nerve electrode placement. A linear incision through the skin and platysma muscle was made one centimeter above the clavicle extending cranially and parallel to the anterior border of the sternocleidomastoid muscle. This muscle was retracted laterally to expose the carotid sheath. The omohyoid muscle was then dissected from the carotid sheath and retracted cranially. The carotid sheath was then opened, and the vagus nerve was exposed between the common carotid artery and jugular vein. At this level, the vagus nerve was transected and exposed to follow it caudally towards any and all branches leading to the heart, lungs and larynx including the superior and inferior cardiac branches, pulmonary branches and the recurrent laryngeal branch. The vagus nerve was sampled below the last pulmonary branch and 2 cm cranial to the average VNS cuff placement. Each sample was approximately 25 cm in length from the upper cervical level to beyond the pulmonary branches at the lower thoracic level. Sutures were placed around the vagus nerve prior to any branching region (i.e., the region where a branch leaves the main vagal trunk) to mark the branching point. Nerves were then placed in neutral buffered formalin (10%) for fixation, labelled accordingly and transported to UCL. All ten nerves were prepared for and scanned, all ten had histology and IHC performed, and five of which had their fascicles segmented.

### Pre-processing and staining

Pre-processing and staining were performed in the same way as the pig vagus nerves in (24). After fixation, nerve samples were measured, sutures of 1 cm length were superglued to the vagal trunk in 4 cm intervals, and nerves cut into 4 cm lengths at the level of suture placement leaving half of the suture on the end of each section as a marker for subsequent co-registration. Two to three sections were placed into a tube of 50 ml Lugol’s solution (total iodine 1%; 0.74% KI, 0.37% I) (Sigma Aldrich L6141) for five days (120 hours) prior to scanning to achieve maximum contrast between fascicles and the rest of the nerve tissue. On the day of the microCT scan, the nerve was removed from the tube and blotted dry on paper towel to remove any excess Lugol’s solution. The nerve sections were placed next to each other onto a piece of cling film (10 cm x 5 cm) (Tesco, United Kingdom) in order from cranial to caudal along the length of the nerve, with cranial ends at the top, and sealed with another piece of cling film to retain moisture during the scan as to avoid shrinkage of the nerve tissue. The sealed nerve samples were rolled around a cylinder of sponge (0.5 cm D x 4.5 cm) and wrapped in another two layers of cling film to form a tightly wound cylinder with a diameter of ∼1.5 cm to fit within the field of view at the required resolution. The wrapped cylinder was placed inside a 3D-printed mount filled with sponge around the edges, ensuring a tight fit and the ends sealed with tape (Transpore™, 3M, United Kingdom).

### MicroCT scanning and reconstruction

A microCT scanner (Nikon XT H 225, Nikon Metrology, Tring, UK) was homed and then conditioned at 200 kVp for 10 minutes before scanning and the target changed to molybdenum. All nerves were scanned with the following scanning parameters: 35 kVp energy, 120 µA current, 7 W power, an exposure of 0.25 fps, optimized projections, and a resolution with isotropic voxel size of 7 µm. Scans were reconstructed in CT Pro 3D (Nikon’s software for reconstructing CT data generated by Nikon Metrology, Tring, UK). Centre of rotation was calculated manually with dual slice selection. Beam hardening correction was performed with a preset of 2 and coefficient of 0.0. The reconstructions were saved as 16-bit volumes and triple TIFF 16-bit image stack files allowing for subsequent image analysis and segmentation in various software.

### Image analysis, segmentation and tracing

This was performed in 5 nerves: 3 left and 2 right. Reconstructed microCT scan images were analyzed in ImageJ (25) in the XY plane to view the cross-section of the nerve. The vertical alignment of the nerve was positioned so that the cross-sectional plane was viewed in the XY stack and the longitudinal plane in the XZ and YZ stacks. This allowed for validation of the scanning protocol, direction of the nerve, and visual analysis of the quality of the image and the distinguishability of the soft tissues – specifically the identification of the fascicles known to exist within the nerve. AVI files were created from ImageJ to enable stack slice evaluation, identification of suture positions and branching locations of the vagus nerve, and as a reference during segmentation. Image stacks (XY plane along the Z-axis) were loaded into Vesselucida 360 (Version 2021.1.3, MBF Bioscience LLC, Williston, VT USA) and image histograms adjusted to optimize visualization of the fascicles when required. Fascicles of the three target organs/functions (namely, cardiac, recurrent laryngeal and pulmonary fascicular groups) were segmented from the rest of the nerve using Vessel mode from the Trace tools. Starting from identification within branches of the vagus nerve, the fascicles were traced through every slice of each scan proceeding proximally up the length of the nerve to the cervical region at the level of cuff placement. Points adding to the path of the vessel trace were place every 50 to 100 sections and the thickness of the vessel trace was altered to match the diameter of the fascicle being traced throughout its length. If fascicles merged with or split into others at a higher frequency, points of the vessel trace were placed at smaller intervals to ensure accurate tracing. The vessel trace was labelled according to the organ from which the fascicle originated. If the fascicle split into two or more fascicles, a bifurcating or trifurcating node was inserted and each branch exiting the node traced proximally up the nerve until either the fascicle merged with another fascicle, or another node needed to be created for splitting fascicles. If fascicles merged into one another, the vessel trace was ended and another trace started that was labelled as containing the respective merged fascicle types and therefore containing fibers from different target organs (i.e., pulmonary and recurrent laryngeal merged fascicle). To continue tracing across cut regions of the nerve, the superglued suture markers and distinct physiological regions or landmarks were used to align the proximal and distal ends of the cut nerves and tracing continued. For visualization of the fully traced nerve, the four overlapping scans were stitched together by aligning the overlapping regions, excluding duplicate images, and forming one large stack of the segmentation data files. Subsequently, this stack was viewed in the 3D Environment where the traces are visualized in 3D displaying the thickness, path and color (specified when labelling the fascicle traces) of the fascicles throughout the length of the scan.

### Histology and IHC for morphology analysis and microCT segmentation validation

Histology and immunohistochemistry (IHC) are useful for identifying nerve anatomical features like nerve diameter, fascicle count and diameter, fiber count, and fiber diameter/type. IHC can further characterize nerves based on neurochemical/microtubule characteristics, distinguishing different fiber types. Trichrome staining uses three dyes to stain collagen, keratin, muscle fibers, cytoplasm, nuclei, and bone (26). By immunostaining with neurofilament (NF) and myelin basic protein (MBP) antibodies on one slide, accurate detection of nerve fiber sizes is possible, visualizing intermediate filaments (27) and myelin sheaths (28) and thereby identifying both myelinated and non-myelinated fibers.

Subsequent to scanning, stained nerves were placed back into neutral buffered formalin for at least a week which allowed for the Lugol’s solution to be soaked out. The formalin was refreshed weekly prior to histology. Histology and IHC was performed on all ten nerves. Segments of nerve 0.5 cm in length, were cut from the human vagus nerves at the level of cuff placement, 2 cm distal to cuff placement, and every 4 cm thereafter. In addition, segments were cut prior to branching and from 0.5 to 1 cm into the branches of the vagus nerve. This was a total of 6 to 7 segments from along the trunk at regular intervals, 3 to 4 segments from above the branching regions, and 3 to 5 segments from the branches of the vagus: this was a total of 9 to 11 segments from the trunk, and a combined total of 12 to 16 segments for histology and IHC per nerve. Segments from the main vagal trunk were placed into a cassette with the order and placement of the segments noted. In another cassette, the segments prior to branching region were placed in order in one row, and those from the branches were placed in order in another row. These nerve samples were embedded in paraffin and six sequential sections cut every 4 µm from each block; two were placed on uncoated slides and three on coated slides. One section of each block on an uncoated slide was stained with Hematoxylin and Eosin (H&E, a routine stain used to demonstrate the general morphology of tissue) (29), the second section on a coated slide stained with Trichrome, and the third section on a coated slide immunostained for both NF (anti-neurofilament heavy polypeptide antibody ab8135 1:1000) and MBP (anti-myelin basic protein antibody ab7349 1:4000) with a secondary biotin labeled antibody (donkey anti-rabbit IgG H&L (Biotin) ab6801 1:1000) (See Supplementary Information). The remaining two sections, one coated and one uncoated, were extra. For H&E staining, a Gemini AS Automated Slide Stainer (Epredia, Portsmouth, New Hampshire, USA) was used. A Leica BOND RX (Leica Biosystems, Milton Keynes, United Kingdom) machine was used for automated IHC. All slides were imaged with light microscopy using a NanaZoomer S360 (Hamamatsu Photonics, Hamamatsu City, Japan).

Identification of histopathological features was performed, and the images of the histology and immunohistochemistry cross-sections were then compared to the corresponding slice in the microCT scan of the same nerve for comparison and validation. The presence and number of fascicles visualized in the golden standard of histology was compared to those identified during segmentation of microCT scans to confirm that what was segmented was correct as describe in Figure S5 and S6 in (24). In addition, fiber sizes within organ-specific fascicles were observed.

### Nerve analysis

NDP.view 2 (Hamamatsu Photonics, Hamamatsu City, Japan) was used to view the images of the histology and IHC cross-sections on the multiple slides. Using NDP.view 2 annotation tools, namely “ruler” and “freehand region”, the diameters (both across the short and long length of the nonuniform circles), area and circumferences were measured for each nerve at the mid-cervical level (Supplementary Table 1). These were then averaged for the ten nerves, and for the left and right nerves, respectively. The number of fascicles (and fascicle bundles – defined as the number of fascicles separated by thin layer of perineurium from but within the same thicker layer of perineurium as other fascicles) were counted at regular intervals along the trunk of the nerve and within the branches for the ten human vagus nerves from the histology images. Specifically, the Trichrome slides were used for this for clear distinguishability of the fascicles from the rest of the nerve tissue. In the cases where a nerve cross-section was not visible on the slides, H&E slides were used. This was compared between nerves (Supplementary Tables 2 and 3). The distance between branches was calculated for all dissected nerves and averaged to provide general gross vagal anatomical information and compared between individuals (Supplementary Table 4). Additionally, the cervical vagus nerve average diameters, areas and fascicle counts were compared to histological studies (Table 1) (4,30–33).

**Table 1.**
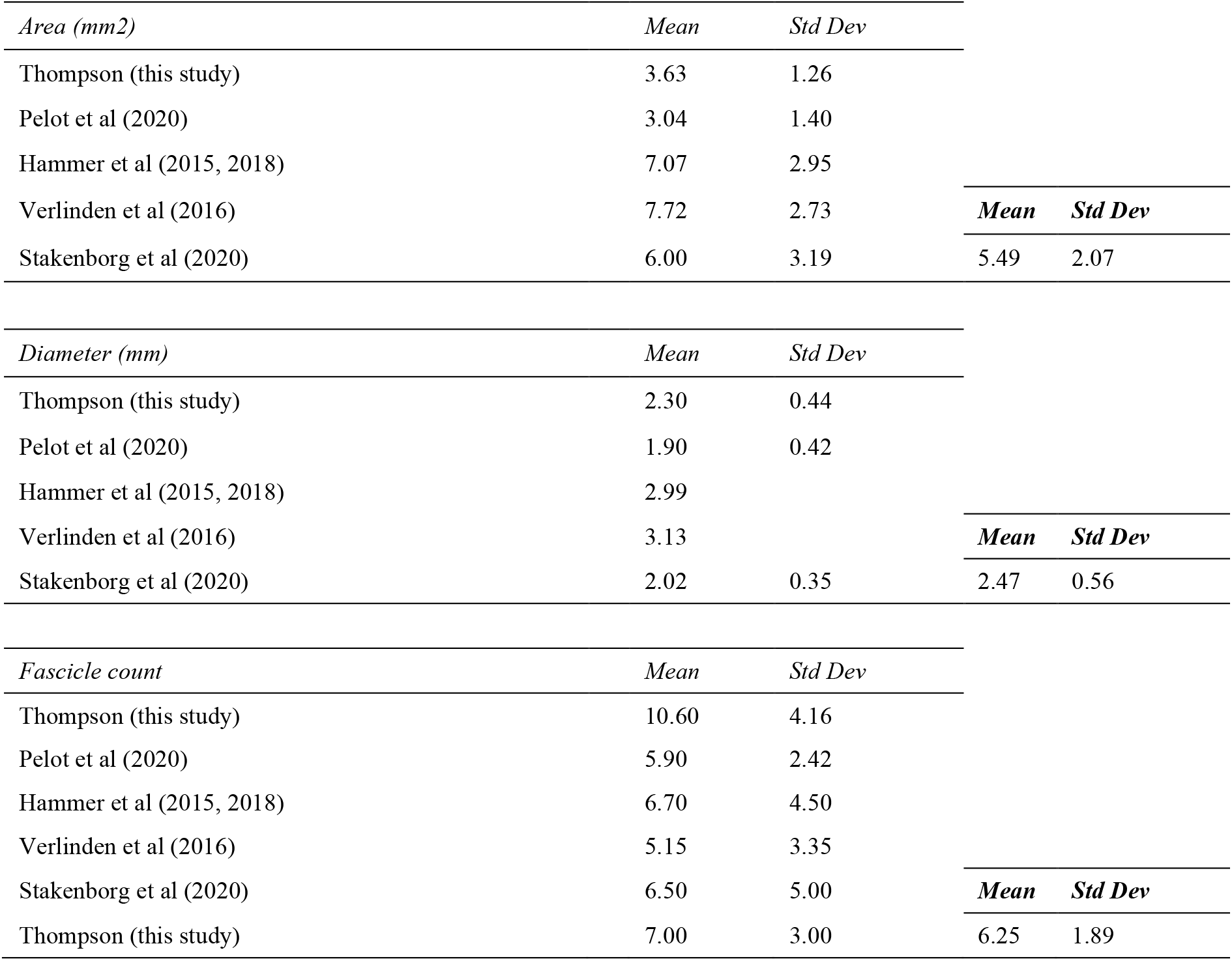
Cervical human vagus nerve morphology compared between studies.

| *Area (mm<sup>2</sup>)* | *Mean* | *Std Dev* |  |  |
| --- | --- | --- | --- | --- |
| Thompson (this study) | 3.63 | 1.26 |  |  |
| Pelot et al (2020) | 3.04 | 1.40 |  |  |
| Hammer et al (2015, 2018) | 7.07 | 2.95 |  |  |
| Verlinden et al (2016) | 7.72 | 2.73 | ***Mean*** | ***Std Dev*** |
| Stakenborg et al (2020) | 6.00 | 3.19 | 5.49 | 2.07 |

| *Diameter (mm)* | *Mean* | *Std Dev* |  |  |
| --- | --- | --- | --- | --- |
| Thompson (this study) | 2.30 | 0.44 |  |  |
| Pelot et al (2020) | 1.90 | 0.42 |  |  |
| Hammer et al (2015, 2018) | 2.99 |  |  |  |
| Verlinden et al (2016) | 3.13 |  | ***Mean*** | ***Std Dev*** |
| Stakenborg et al (2020) | 2.02 | 0.35 | 2.47 | 0.56 |

| *Fascicle count* | *Mean* | *Std Dev* |  |  |
| --- | --- | --- | --- | --- |
| Thompson (this study) | 10.60 | 4.16 |  |  |
| Pelot et al (2020) | 5.90 | 2.42 |  |  |
| Hammer et al (2015, 2018) | 6.70 | 4.50 |  |  |
| Verlinden et al (2016) | 5.15 | 3.35 |  |  |
| Stakenborg et al (2020) | 6.50 | 5.00 | ***Mean*** | ***Std Dev*** |
| Thompson (this study) | 7.00 | 3.00 | 6.25 | 1.89 |

## Results

### MicroCT scanning and fascicle tracing of human vagus nerves

The organization of the cardiac, pulmonary and recurrent laryngeal was present directly after all the branches entered the vagus nerve in all nerves (Figure 1-7) (n=3). Detailed morphology of the nerve and its branches can be seen in the ‘Analysis of nerve morphology’ section. In the left vagus nerves, approximately 1-1.5 cm up the nerve, cardiac fascicles merged with some pulmonary fascicles. The remaining cardiac fascicles merged after 2.5 cm. The recurrent laryngeal fascicles remained on one half of the nerve for at least 2-3 cm and then eventually all fascicles of the recurrent laryngeal merged by 6-7 cm proximally from the branch entry point. In the right vagus nerves, the inferior cardiac not only predominantly merged with the pulmonary fascicles (forming a common, pink fascicle, Figure 1-7) as in the left vagus nerves, but in both right vagus nerve samples segmented, the inferior cardiac fascicles merged with a recurrent laryngeal fascicle (forming a common, yellow fascicle, Figure 6 and 7) at 1.7 and 2.1cm, respectively, from point of entry in the nerve till merger point. This difference of merging pattern may be attributed to the differing branching point of recurrent laryngeal branch between left and right vagi. At the point where all three organ-specific fascicle types had merged, microCT has a limitation of not allowing for the visualization and thus, the tracing of the fibers of the nerves. Therefore, the organization of the organ-specific fibers within these fascicles remains unknown. The vagus nerve structure became plexiform after all the three organ-specific fascicles merged together (Figure 8). The human vagus nerve appeared to merge or split at an approximate rate of one to two fascicles every 0.5 mm. Further up the nerve, below the cervical level, a superior cardiac branch entered the vagus nerve and remained separated from the other fascicles for a short length (2-3 cm) which is just below VNS stimulator placement. In summary, so far, organization and separation between the organ-specific fascicular groups exists at approximately clavicular level and the superior cardiac fascicles are separate from the other vagal fascicles at a position close to VNS cuff placement.

**Figure 1.**
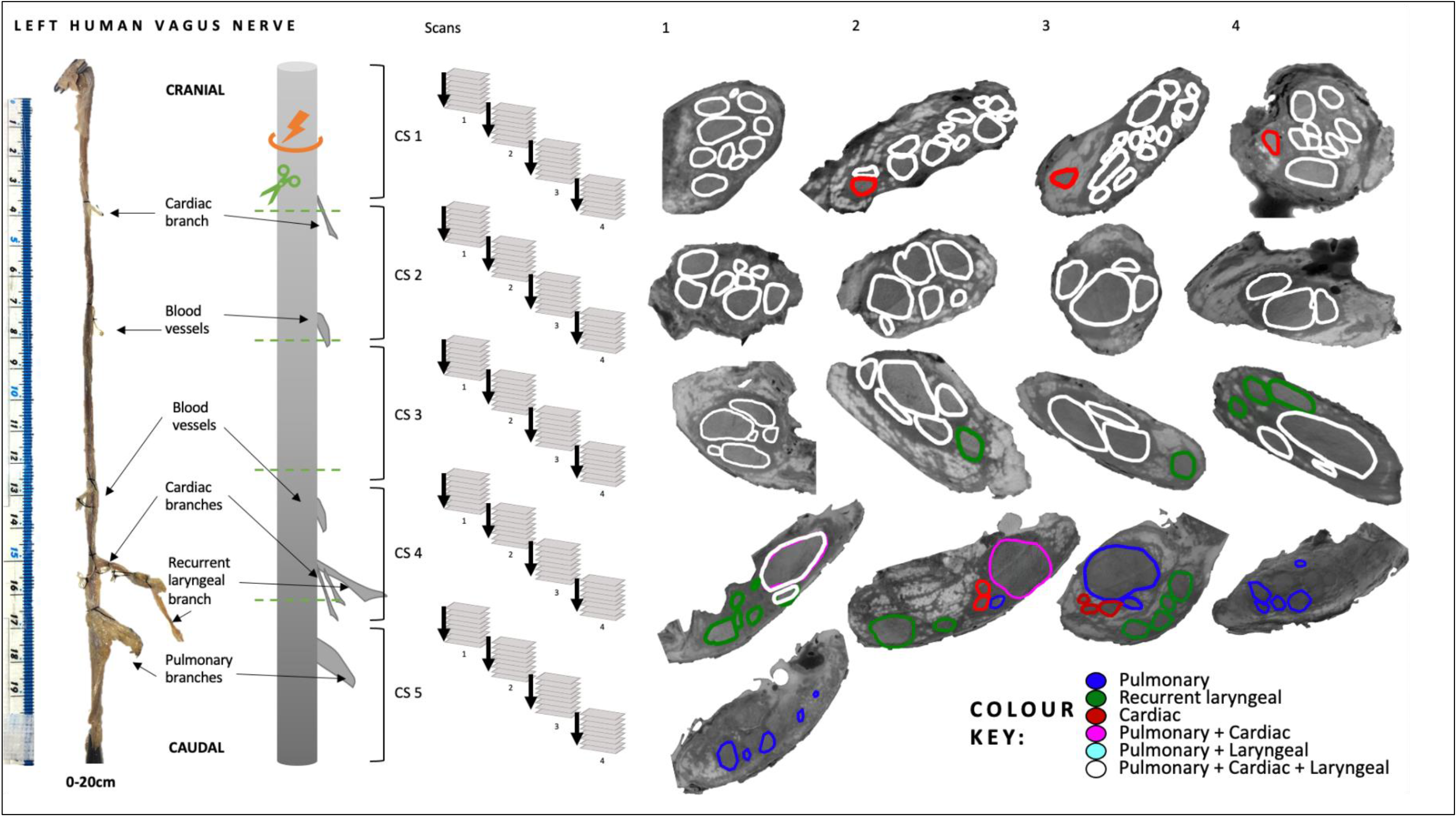
Human vagus nerve preparation, scanning setup and segmentation at intervals. The vagus nerve samples with branches marked with sutures around the trunk were cut (green dashed line) into 4 cm segments prior to staining identified here as cross-sections (CS) 1 to 5. The segments were bundled together and scanned with four overlapping scans. This figure illustrates how, when viewing one CS, four scans were required to image its full length. CSs from the top of each scan, proceeding proximally up the nerve from CS5 to CS1, are displayed here for one nerve example with labelled, segmented fascicles to illustrate the progression of fascicle organization and movement within the nerve.

**Figure 2.**
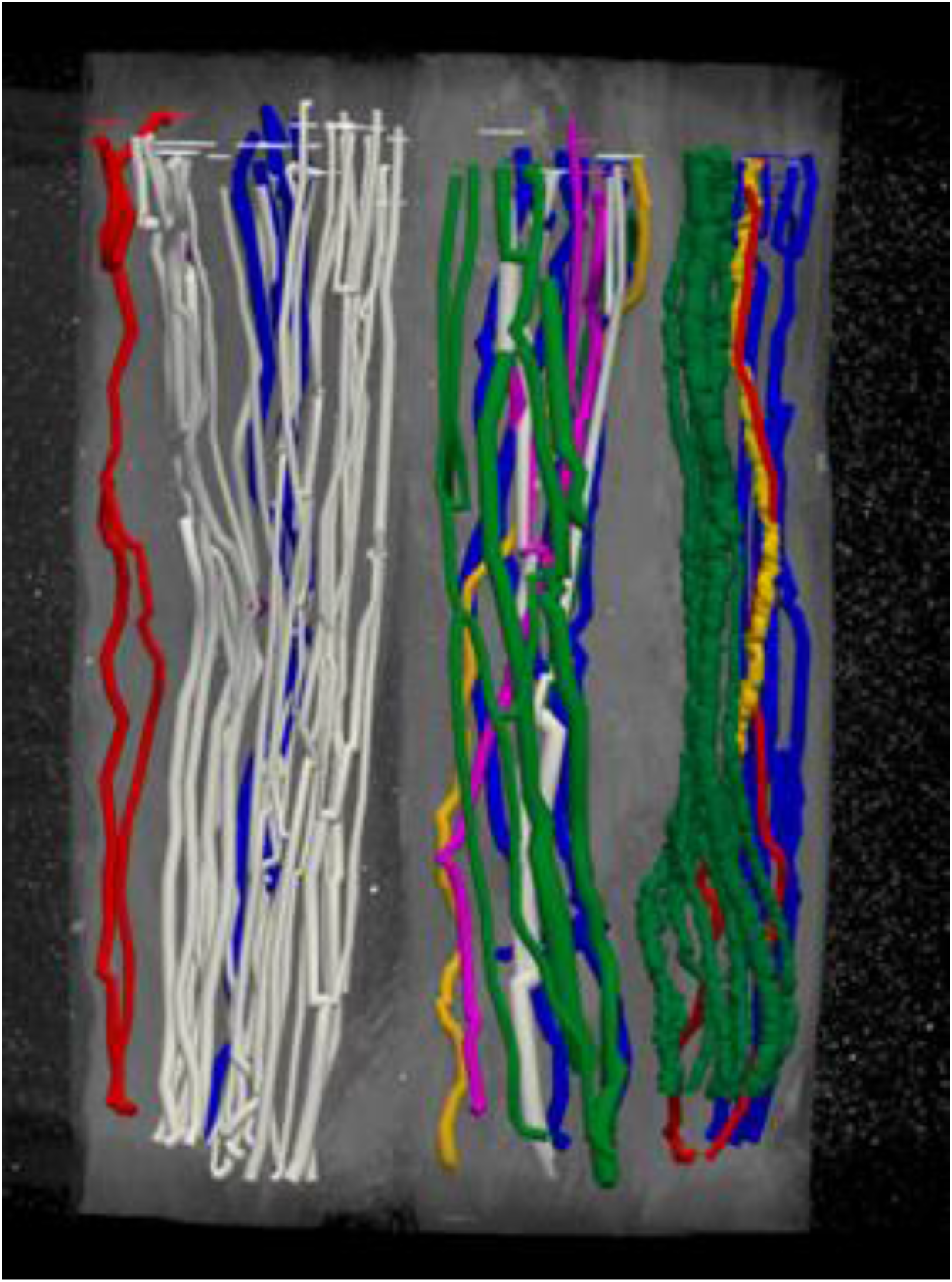
Segmentation through multiple cross-sections in a scan. An example of scan with multiple segments/cross-sections of nerve. The fascicles have been traced through the cross-sections through the overlapping scans.

**Figure 3.**
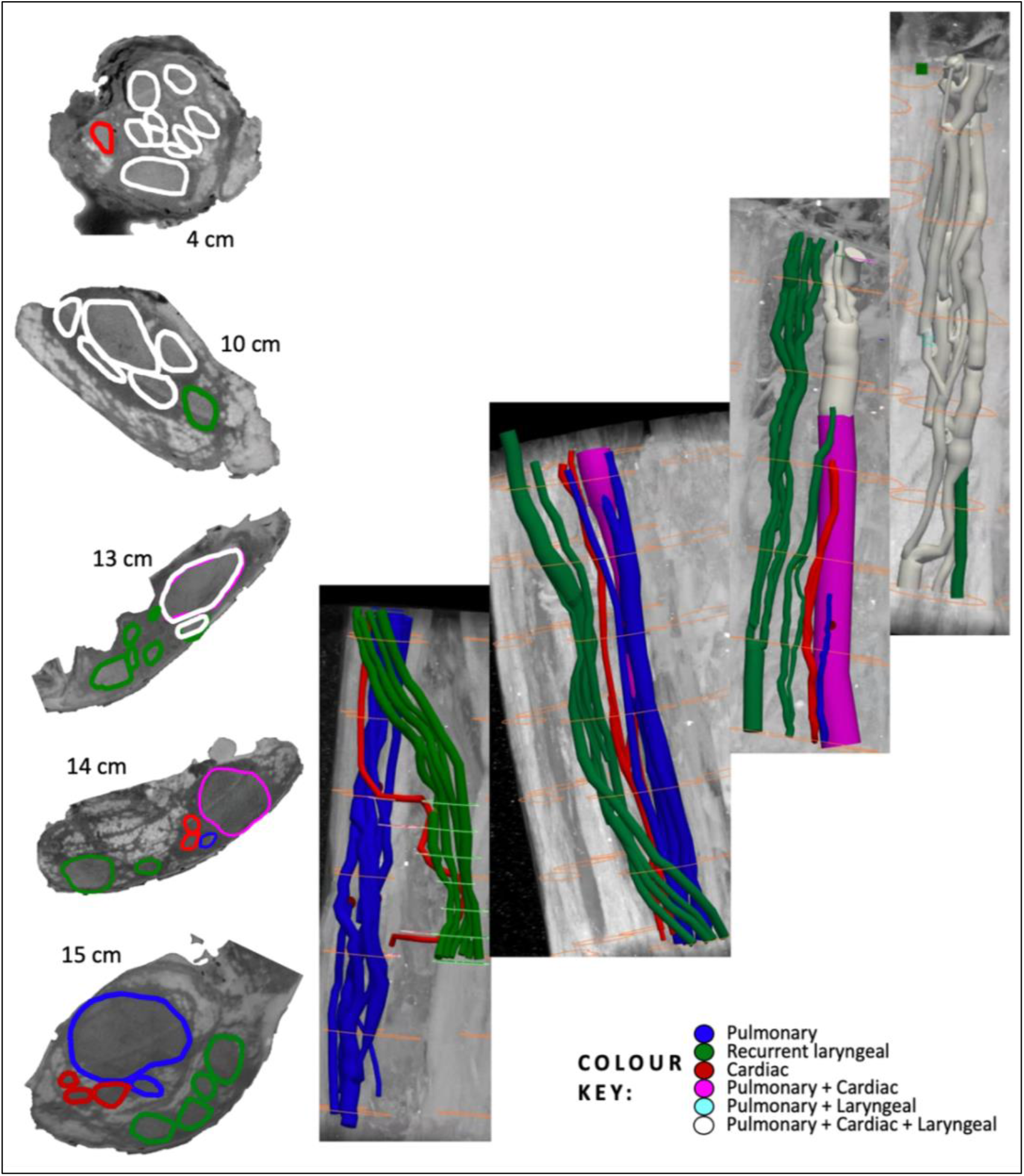
Human vagus nerve segmentation 1 - left. Segmentation of fascicles in 3D from caudal (bottom/left) to cranial (top/right) for one left vagus nerve to the point at which most fascicles have merged (white) approximately 4 cm from all three branches entering the vagus nerve (excluding superior cardiac branches). The CS of the nerve at specified intervals from cervical level are displayed on the left. The cardiac fascicle present in the first CS is the superior cardiac fascicle(s). In the second segmentation figure from the left, after the recurrent laryngeal and cardiac fascicles enter the vagus nerve from their branches (right), the cardiac branches merge with the pulmonary fascicles shortly thereafter (left, pink).

**Figure 4.**
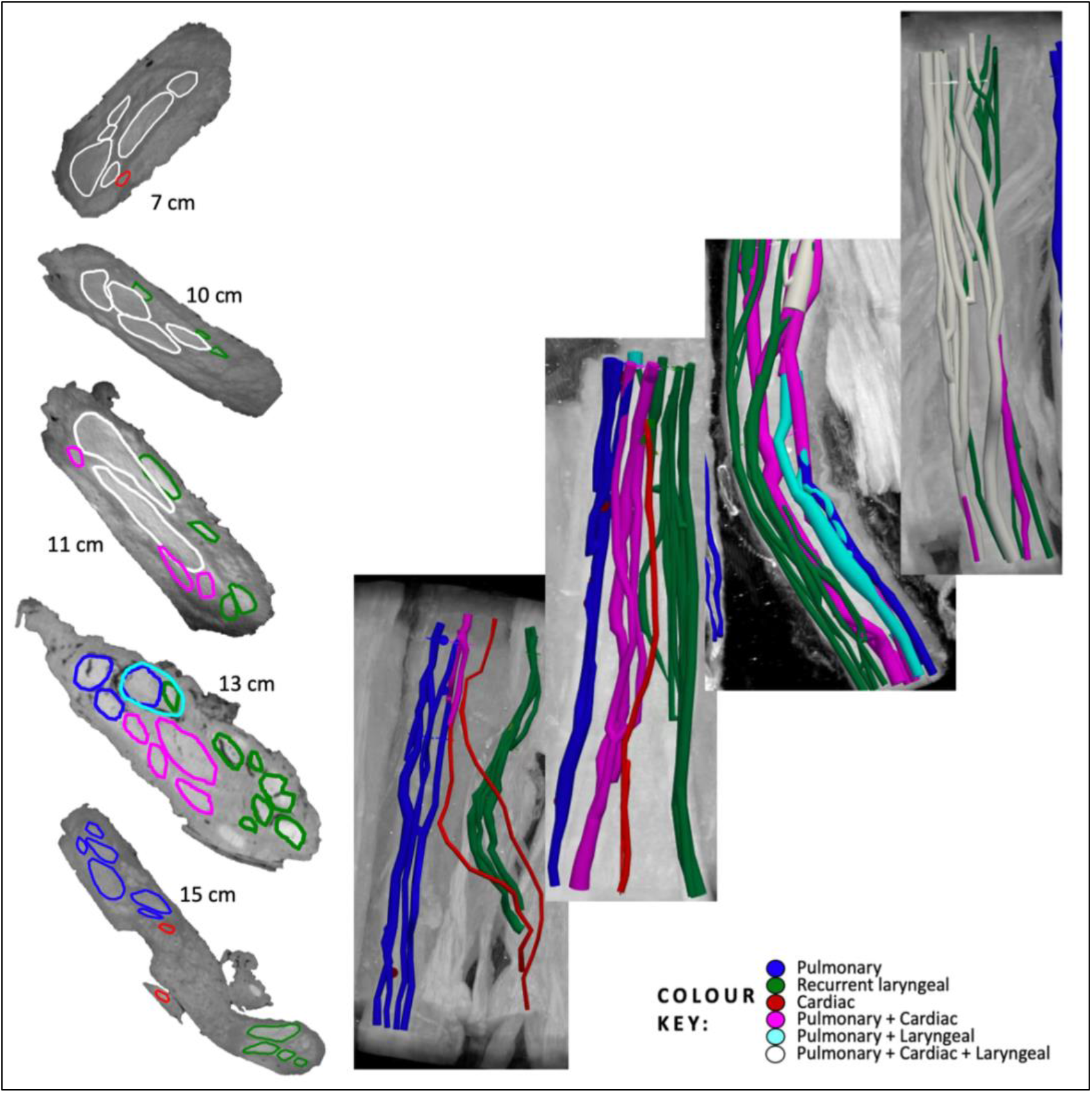
Human vagus nerve segmentation 2 - left. Segmentation of fascicles in 3D from caudal (bottom/left) to cranial (top/right) for a second left vagus nerve to the point at which most fascicles have merged (white) approximately 4 cm from all three branches entering the vagus nerve (excluding superior cardiac branches). The CS of the nerve at specified intervals from cervical level are displayed on the left. The cardiac fascicle present in the first CS is the superior cardiac fascicle(s).

**Figure 5.**
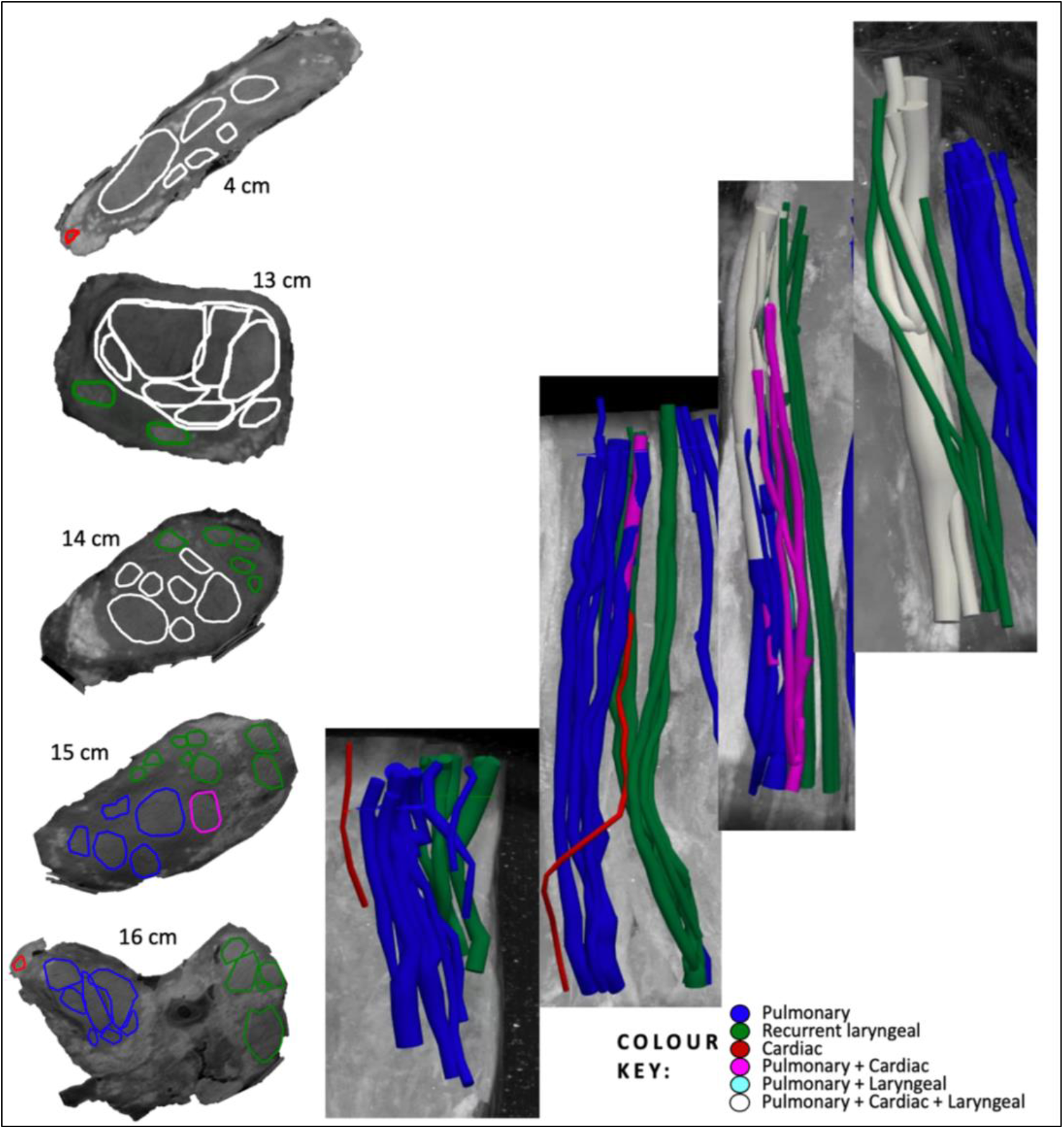
Human vagus nerve segmentation 3 - left. Segmentation of fascicles in 3D from caudal (bottom/left) to cranial (top/right) for a third left vagus nerve to the point at which most fascicles have merged (white) approximately 4 cm from all three branches entering the vagus nerve (excluding superior cardiac branches). The CS of the nerve at specified intervals from cervical level are displayed on the left. The cardiac fascicle present in the first CS is the superior cardiac fascicle(s).

**Figure 6.**
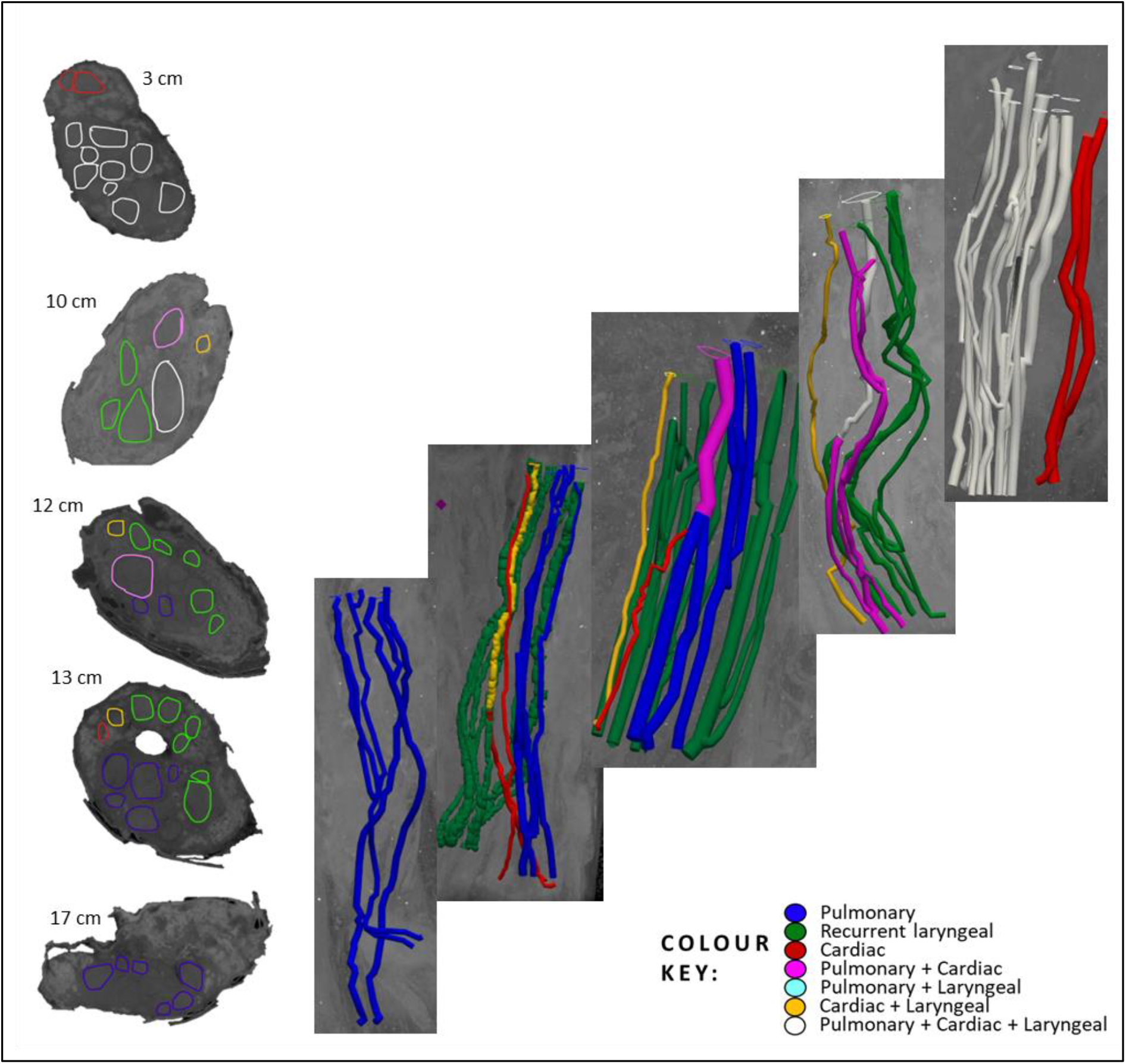
Human vagus nerve segmentation 4 - right. Segmentation of fascicles in 3D from caudal (bottom/left) to cranial (top/right) for a fourth nerve from the right side, to the point at which most fascicles have merged (white) approximately 4 cm from all three branches entering the vagus nerve (excluding superior cardiac branches). The CS of the nerve at specified intervals from cervical level are displayed on the left. The cardiac fascicle present in the first CS is the superior cardiac fascicle(s).

**Figure 7.**
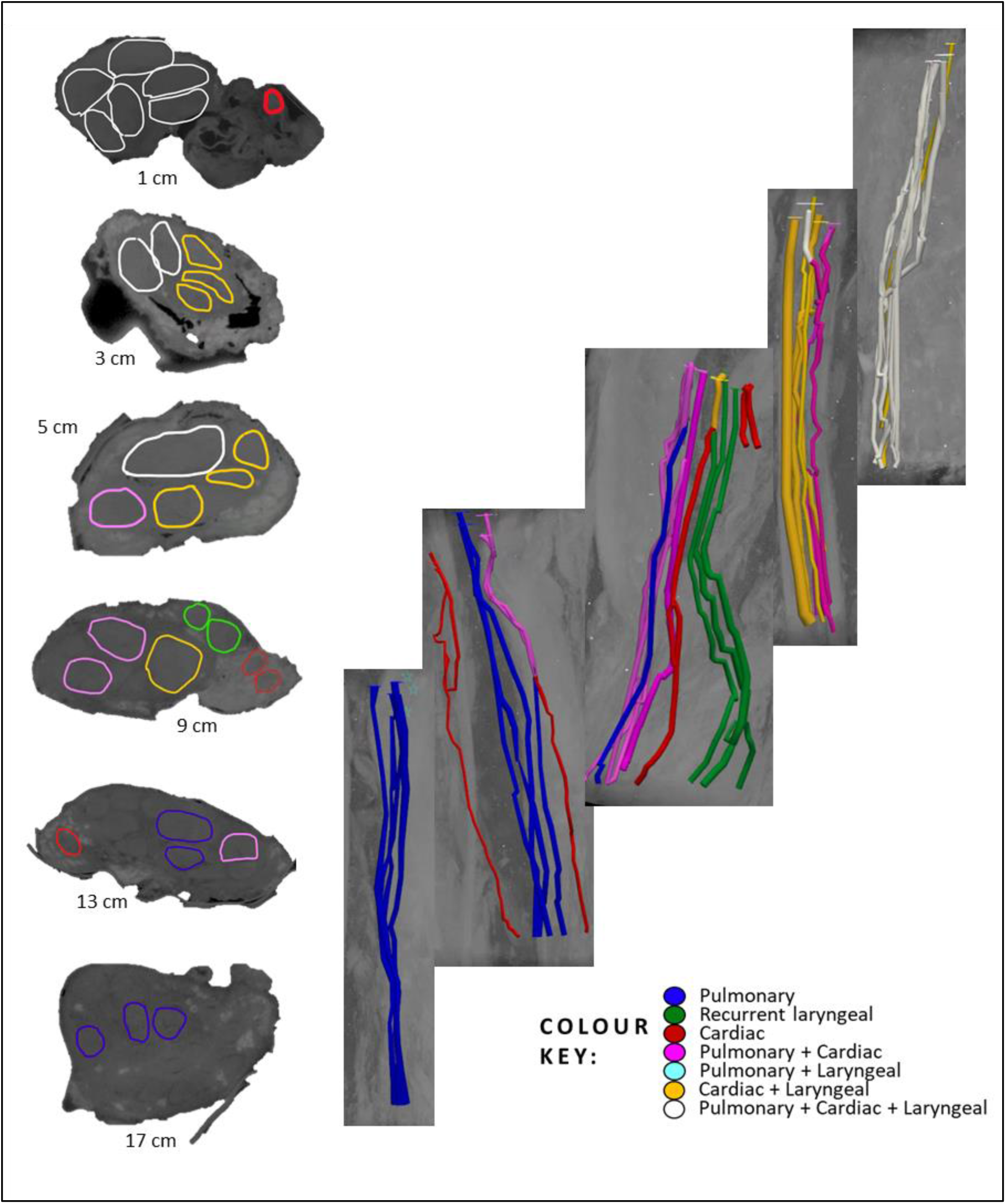
Human vagus nerve segmentation 5 - right. Segmentation of fascicles in 3D from caudal (bottom/left) to cranial (top/right) for a fifth, right vagus nerve to the point at which most fascicles have merged (white) or fascicles containing cardiac and laryngeal fibers (yellow) approximately 4 cm from all three branches entering the vagus nerve (excluding superior cardiac branches). The CS of the nerve at specified intervals from cervical level are displayed on the left. The cardiac fascicle present in the first CS is the superior cardiac fascicle(s).

**Figure 8.**
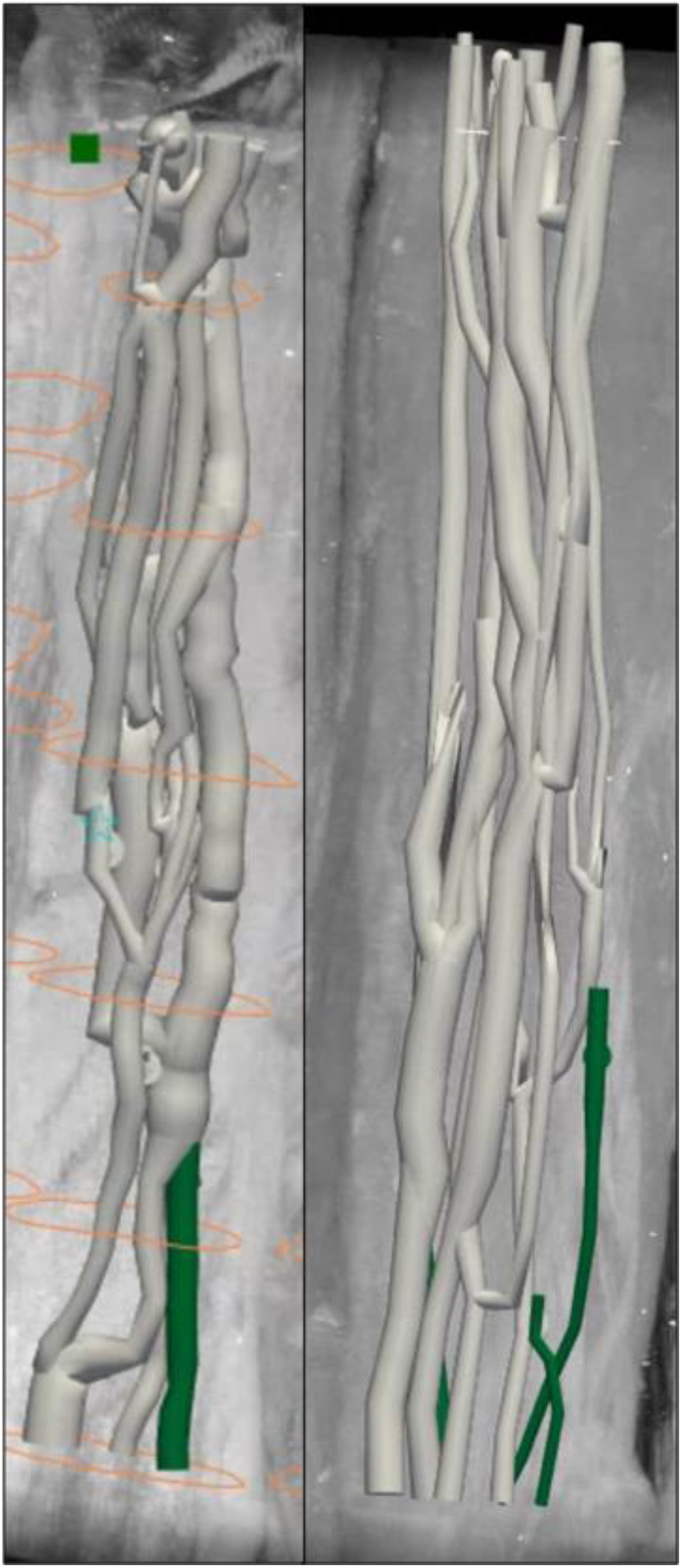
Anastomoses observed in human vagus nerve. Two examples of the frequent anastomosing (merging and splitting) between and plexiform structure of fascicles observed in the human vagus nerve. Once organ-specific fascicles of different types merge together, the fascicular resolution of microCT scans does not allow for the tracing and identification of organ specificity after merging and during these anastomoses; therefore, from this point on, labelled as white “merged” fascicles. For another example, please see <u>Supplementary Video 1</u>.

### Histology and immunohistochemistry

Histology and immunohistochemistry were performed on all nerves including H&E, Trichrome, NF with MBP. An example of all three stains on two example cross-sections can be seen in Figure 9. NF with MBP successfully stained fibers simultaneously, allowing for the visualization of the varying fiber sizes within the cross-sections with ease. Examples of this visualization can be seen in Figure 10 and 11. The large, myelinated, efferent fibers in the recurrent laryngeal fascicles (Figure 10E, 11D) are evident in the cross-sections from the branch as well as in the vagal trunk just proximal to the entry of the fascicles. At the cervical level (Figure 10A, 11A), however, the fibers appear to be mixed with large fibers dispersed around the cross-section and only small clusters of large fibers observed. Despite hypotheses that the clusters of fibers will remain organ-specific, this cannot be deciphered without tracing of the fibers throughout the nerve.

**Figure 9.**
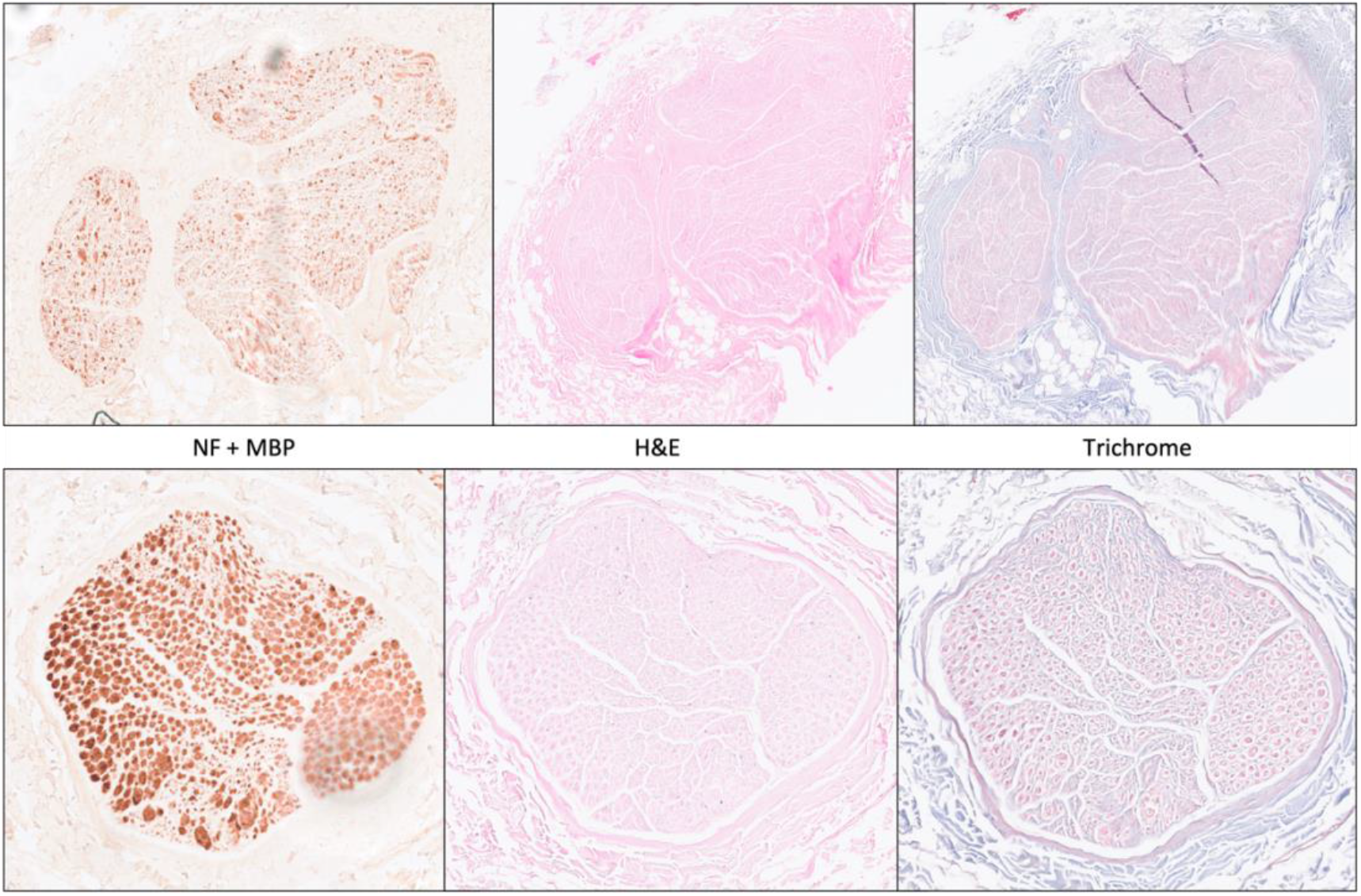
Two examples of the different histological and immunohistochemical stains used. Cross-sections from two nerves with each of the histological or immunohistochemical stains including double staining with neurofilament (NF) and myelin basic protein (MBP), hematoxylin and eosin (H&E), and Trichome staining, from left to right, respectively. This was performed for all 10 nerves at numerous intervals along the nerve.

**Figure 10.**
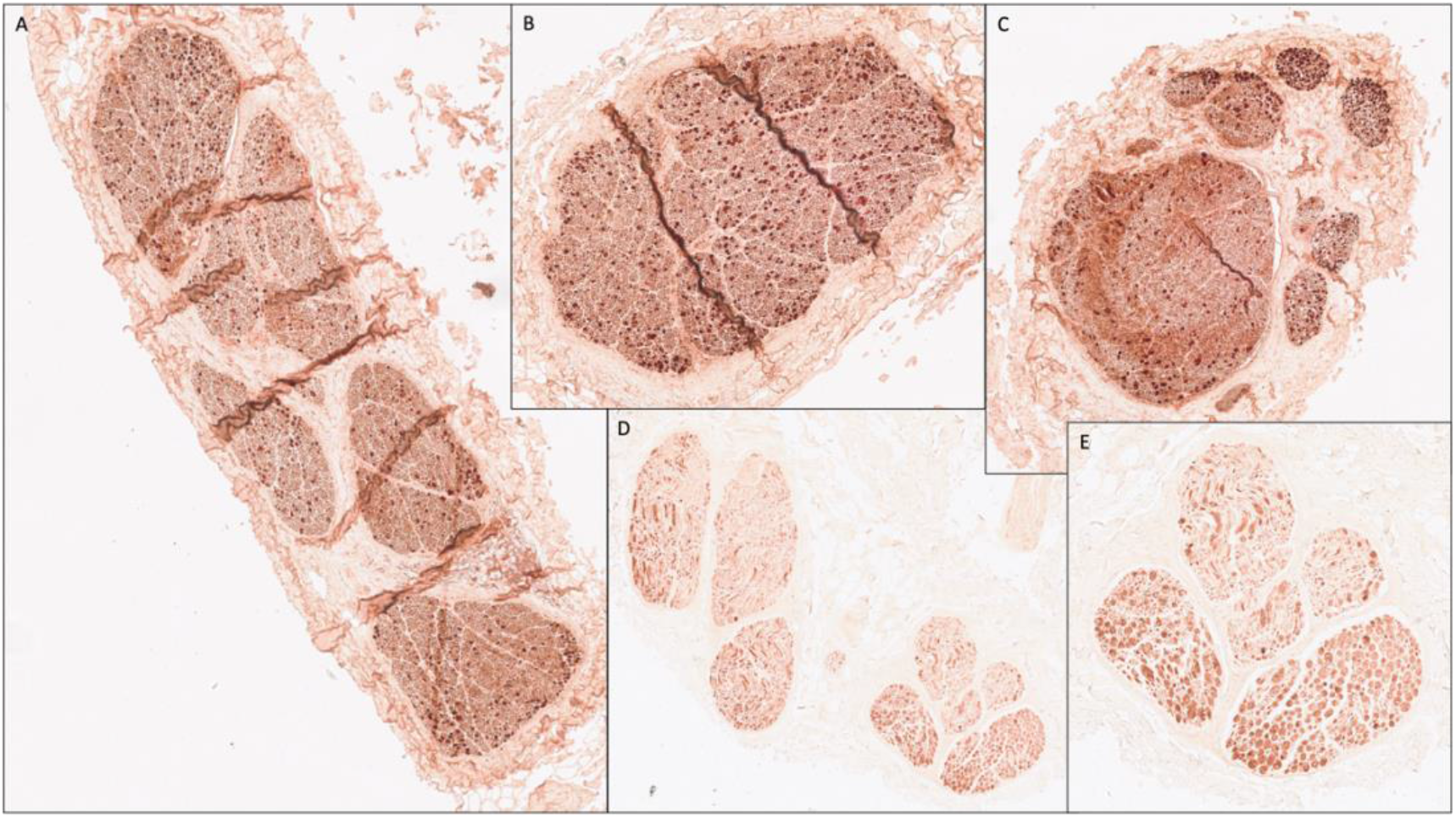
IHC of a human vagus nerve example 1. An example of strong double staining of neurofilament and myelin basic protein with cross-sections at: A. Cervical level; B. 5cm proximal to recurrent laryngeal branching point (one fascicle present); C. Recurrent laryngeal fascicles in vagus nerve trunk prior to branching; D. Recurrent laryngeal branch; E. Recurrent laryngeal fascicles (higher zoom/resolution).

**Figure 11.**
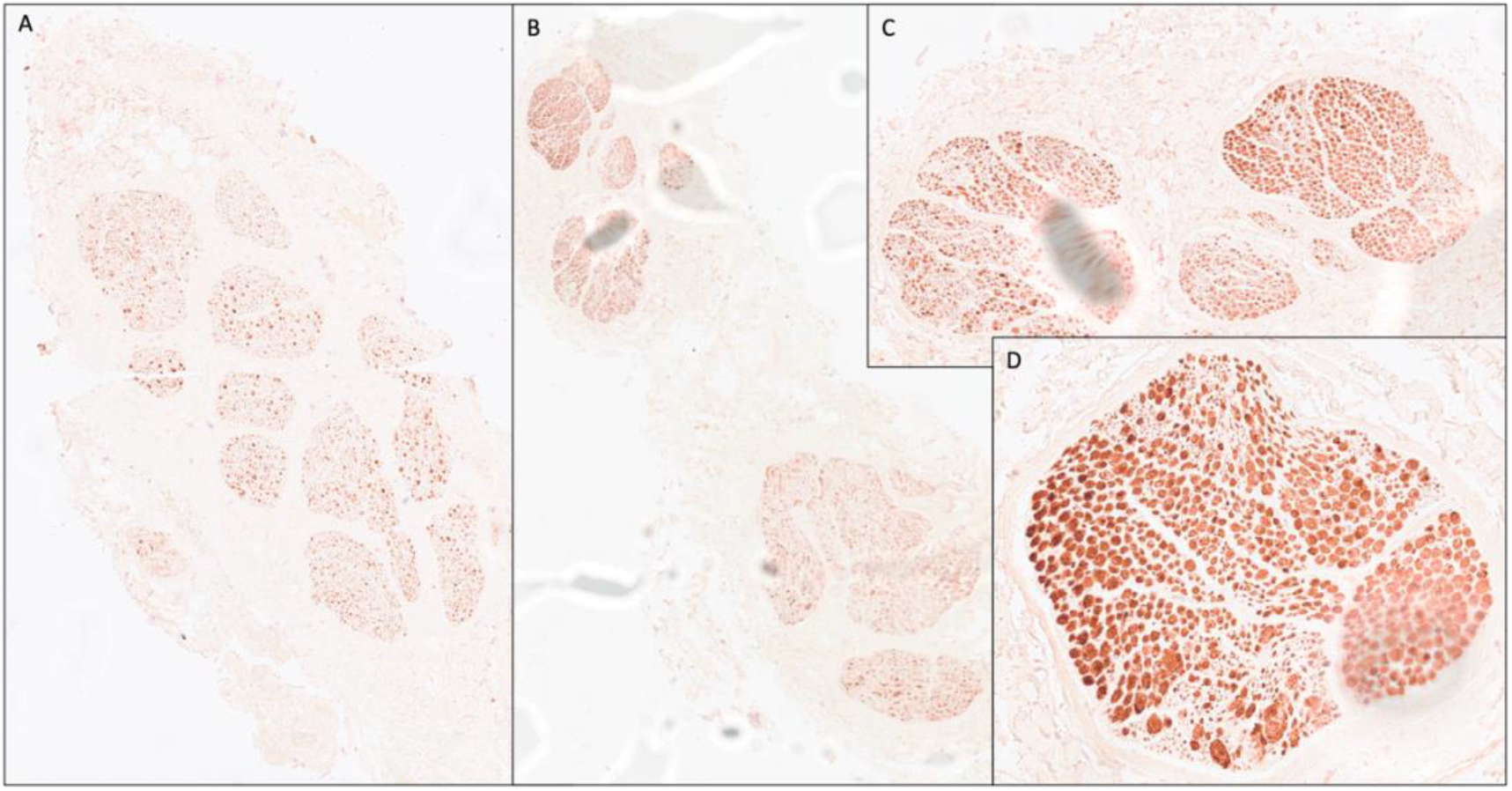
IHC of a human vagus nerve example 2. Another example of double staining of neurofilament and myelin basic protein with cross-sections at: A. Cervical level; B. Recurrent laryngeal fascicles in vagus nerve trunk prior to branching; C. Recurrent laryngeal branch; D. Recurrent laryngeal fascicle (higher zoom/resolution).

Histology and immunohistochemistry allowed for validation of microCT segmentation – confirming the presence of fascicles from that which was segmented from the scans. A nerve example of microCT compared with immunohistochemistry can be seen in Figures 12, 13 and 14. Regions of the stack of XY slices of microCT data from where the histology slice was taken were compared to the histology slice of interest to find the matching slice for accurate cross-validation. For display purposes here, the closest already segmented and labelled cross-section was compared and therefore, there is slight mismatch. Recurrent laryngeal fascicles can be seen to contain more larger, myelinated fibers than the other fascicles in the nerve. The mixed (cardiac and pulmonary) fascicles contained a mix of fiber sizes. The superior cardiac (Figure 14) fascicle also contained a mixture of fiber sizes.

**Figure 12.**
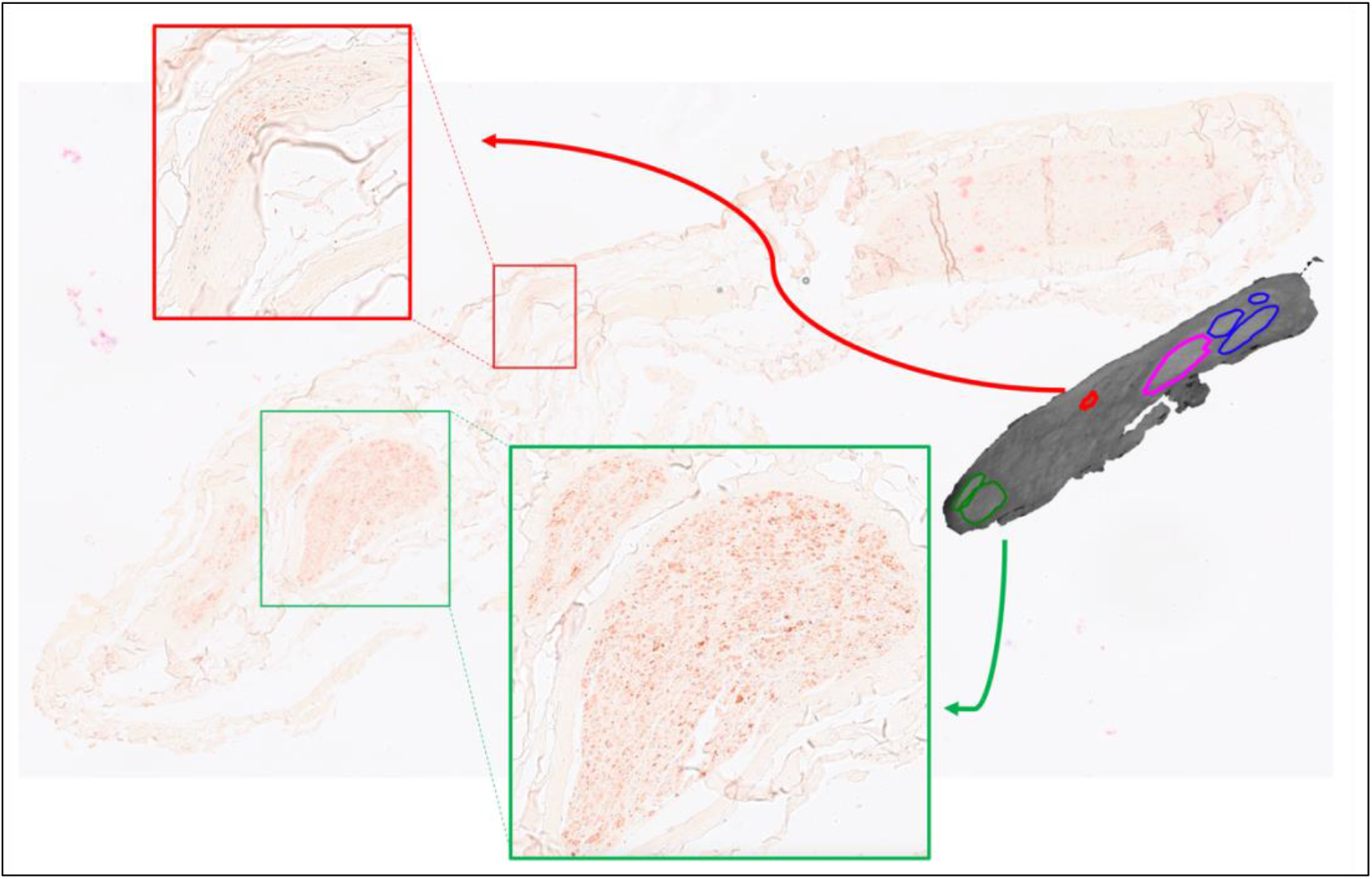
MicroCT with IHC 1. An example from one nerve of IHC compared to microCT 14cm from mid-cervical level with cardiac (red) and recurrent laryngeal (green) fascicles highlighted. Pulmonary (blue) fascicles and a merged fascicle between cardiac and pulmonary (pink) are also visible in the microCT cross-section.

**Figure 13.**
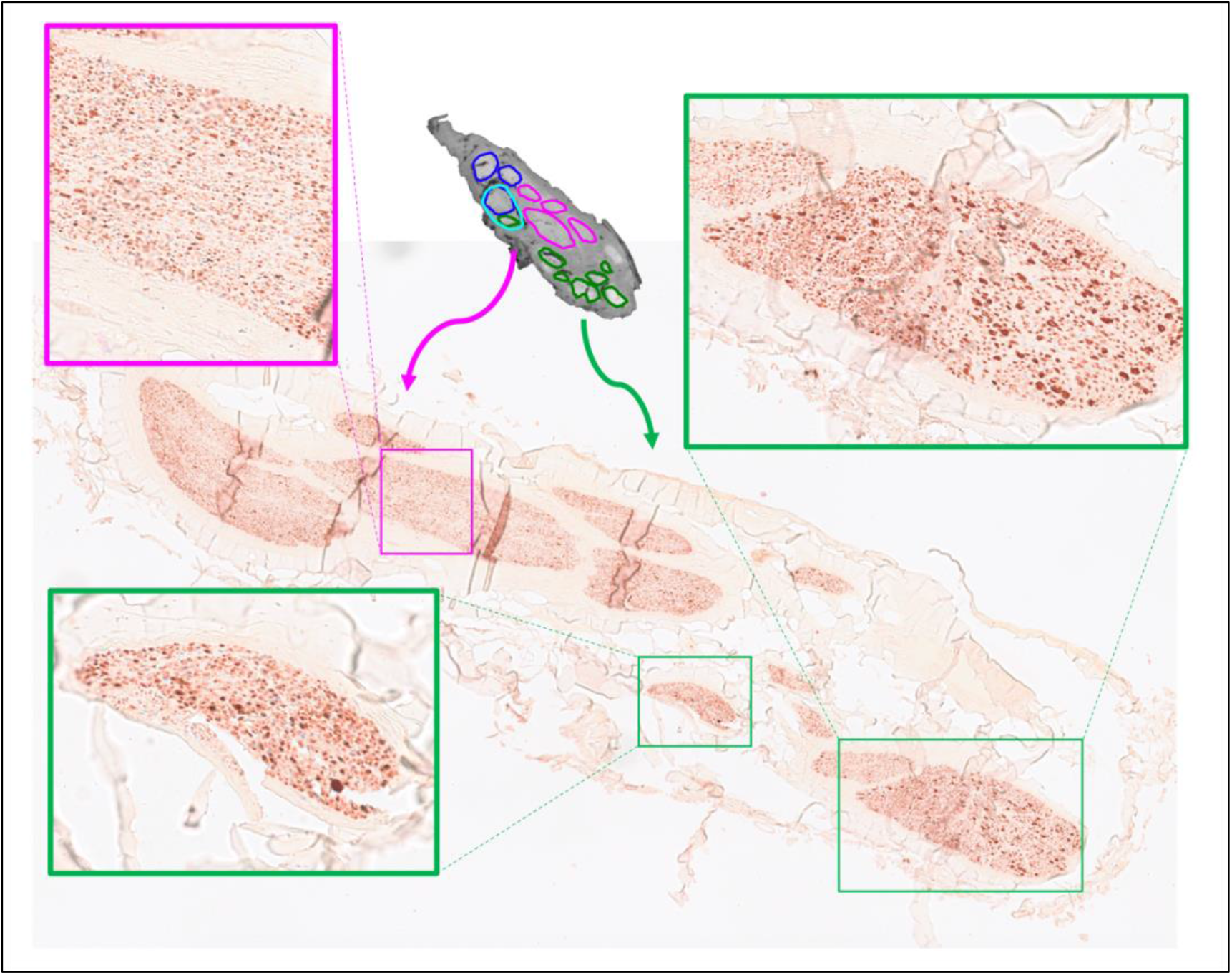
MicroCT with IHC 2. An example from one nerve of IHC compared to microCT 13cm from mid-cervical level with merged between cardiac and pulmonary (pink) and recurrent laryngeal (green) fascicles highlighted. Pulmonary (blue) fascicles are also visible in the microCT cross-section.

**Figure 14.**
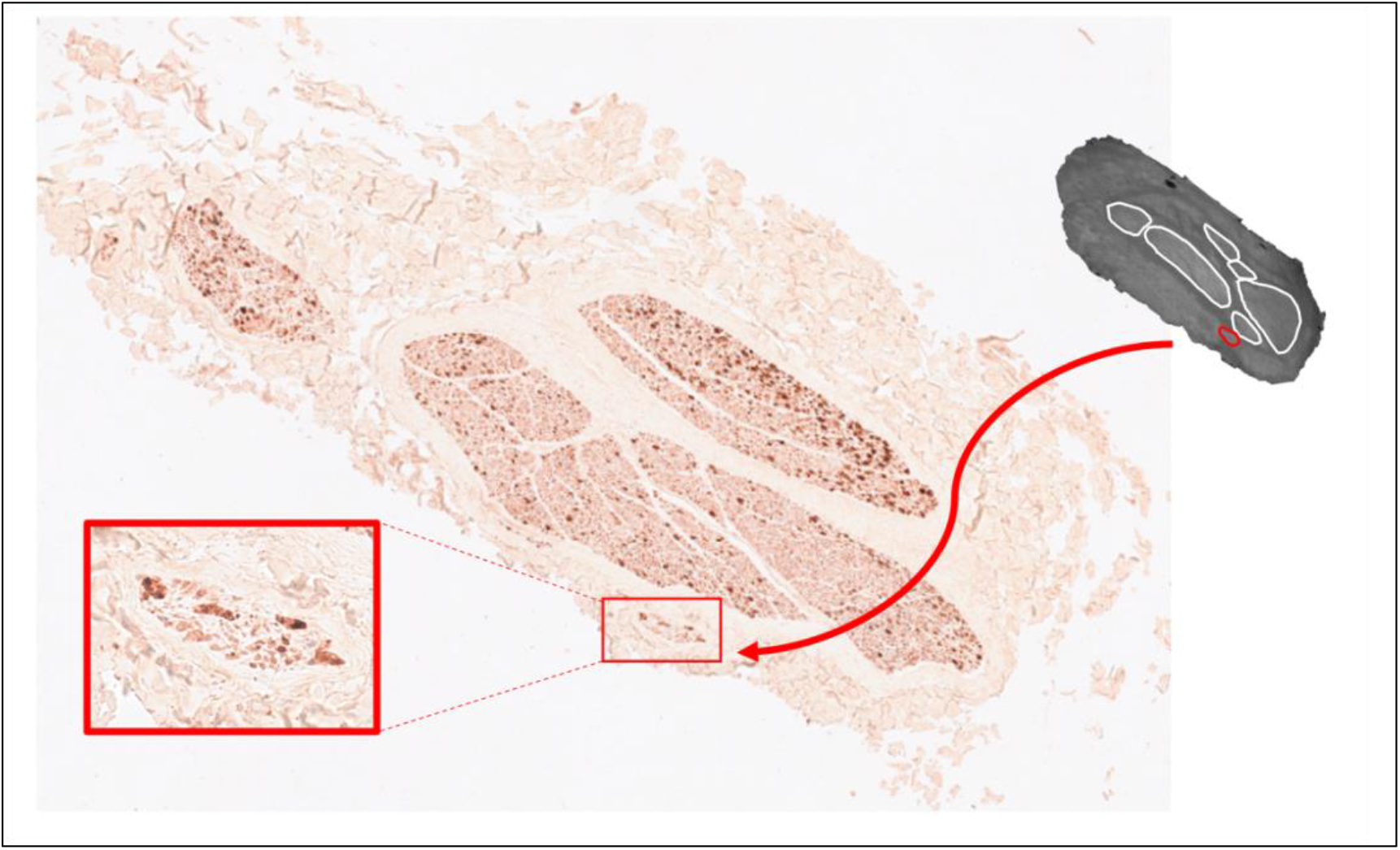
MicroCT with IHC 3. An example from one nerve of IHC compared to microCT 7cm from mid-cervical level with a superior cardiac fascicle (red) highlighted. Merged fascicles between inferior cardiac, recurrent laryngeal and pulmonary (white) are also visible in the microCT cross-section.

### Analysis of nerve morphology and anatomy from microCT and histology

From the morphological data of the nerve, the right human vagus nerves were calculated to be on average 1.22 times larger than the left vagus nerve in terms of its diameters, circumference, and area, with an average area of all 10 nerves of 3.63 mm^2^ ± 1.43, an average area of the left vagus nerves of 3.08 mm^2^ ± 1.77, and an average area of the right vagus nerves of 4.17 mm^2^ ± 0.85. In addition, the right vagus nerves contain on average 1.28 times more fascicles and fascicle bundles than the left with a mean of 10.60 ± 4.12 fascicles for all nerves, 9.60 ± 3.05 left vagus nerve fascicles, and 11.60 ± 5.13 right vagus nerve fascicles (Supplementary Table 1, Supplementary Figure 1). The left and right cervical vagus nerves consisted of 5.00 ± 1.58 and 6.8 ± 2.95 fascicle bundles, respectively.

The number of fascicles present in the cross-sections of the main trunk of the vagus nerve varied when progressing through the nerve at regular intervals (Supplementary Tables 2 and 3). The least number of fascicles was an average of 8.30 ± 3.43 and 8.67 ± 4.5 at 1 cm and 13 cm from the upper cervical level, respectively. At 9 cm from upper cervical level, there was the greatest average number of fascicles across the 10 nerves of 16.20 ± 7.28. The fascicle counts increasing and decreasing when progressing down the length of the nerve illustrate the anastomoses taking place. This coincides with the high standard deviations observed here such as a standard deviation of 10.19 for nerve 3. Averaging all 10 nerves’ mean fascicle counts from regular intervals along the nerves’ length gives a mean fascicle count of 11.85 ± 5.75. This is greater than the mean at cervical level. On average between nerves, the superior cardiac branch contained 3.10 ± 1.45 fascicles, the inferior cardiac branch 2.33 ± 1.06, recurrent laryngeal branch 7.25 ± 1.98, and the pulmonary branch 6.14 ± 2.91 fascicles.

Unlike the pig vagus nerves, the human inferior cardiac branches exit the vagus nerve just above or at the same time with the recurrent laryngeal branches (Supplementary Table 4). The distance from cervical level on average to the inferior cardiac branches and recurrent laryngeal branch was 12.30 cm ± 1.89 and 12.30 cm ± 1.89, and 15.30 cm ± 1.44 and 15.00 cm ± 1.27 for the right and left nerves, respectively. The discrepancy of the branching point of the right and left recurrent laryngeal branches of the vagus nerve is expected. The superior cardiac branch exits the vagus nerve on average 6.38 cm ± 2.43 from the upper cervical level with an average difference of 2.45 cm between the left and right nerves with the branch exiting at 5.15 cm and 7.60 cm, respectively. The pulmonary branches exited the vagus nerve from 18 to 20 cm from the upper cervical level. Four fifths of the right vagus nerves had three pulmonary branches and four fifths of the left vagus nerve has two pulmonary branches which coincides with the number of lobes the lungs consist of on either side. There was an average of 0.95 cm between each pulmonary branch.

Compared to morphological data previously reported in literature, the effective diameters were comparable between studies (Table 1). In addition, the fascicle counts, when looking at fascicle bundles in this work, was comparable between studies too. The individual fascicle count reported here was larger; this may be indicative of the other studies not taking into account thin layers of perineurium. The nerve cross-sectional areas correlated with Pelot et al (2020) but the other three studies reported greater areas of 1.5 times, albeit with overlapping ranges.

## Discussion

In summary, the segmented human vagus nerves show a fascicular, organotopic organization only until clavicular level. Thereafter, the fascicles anastomose and form a plexiform structure within the nerve from clavicular level until cervical level with a frequency of one merge or split every 0.5 mm. The superior cardiac fascicle(s) are still separate from the rest of the merged vagal fascicles at level of cuff placement (n=3, 1 left, 2 right) or just below (n=2 left) still within the mid-cervical level in a readily accessible location for cuff placement. The merging pattern between fascicles of different organ origin was reasonably similar between nerve samples and between sides; however, there was one notable difference between left and right, with the inferior cardiac branch fascicles of the right vagus nerve not only merging predominantly with the pulmonary fascicles, as on the left side, but with one fascicle also merging with a recurrent laryngeal fascicle. This difference of merging pattern between sides may be attributed to the different average location of the recurrent laryngeal branching point on the vagus. On average, at cervical level, the right vagus contained 11.6 and 6.8 and the left 9.6 and 5 fascicles and fascicle bundles, respectively. The cross-sectional areas of the right and left vagus nerves were 4.17 mm^2^ ± 0.85 and 3.08 mm^2^ ± 1.77, respectively. The number of fascicles comprising the vagus nerve at regular intervals varied significantly, increasing and decreasing along its length with only an observed pattern of the most fascicles being present at 9 cm from mid-cervical level on average across ten nerves. Histology allowed for validation of the microCT segmentation and allowed for visualization of fiber size dispersion and clusters present in the nerve. The large fibers observed in recurrent laryngeal fascicles from its branch appear to be dispersed around fascicles of the nerve with only small clusters still remaining at cervical level.

The morphometric data was comparable with previously reported data in terms of diameter, area and fascicle counts (4,30,32,33) (Table 1). The cross-sectional areas reported in three of the studies was greater than those reported here and in Pelot et al (2020). However, the diameters were similar. This may indicate differences in calculations of area between studies. These data together should be used as an informative dataset for human cervical vagus nerve morphology which could inform cuff design, stimulation strategies and cuff placement. Averaging the five studies’ data, the human cervical vagus nerve has an area of 5.49 mm^2^ ± 2.07, a diameter of 2.47 mm ± 0.56, and 6.25 ± 1.89 fascicles.

The frequency of splitting and merging (anastomosing) correlated with that previously quantified over 2 cm of cervical vagus nerve (34). This also corresponds with the large variance in fascicle counts along the length of the nerve at regular intervals observed for all ten nerves. The high degree of fascicular anastomoses along with the variations in key morphological characteristics of fascicles across humans may attribute for the dispersion of fibers observed at cervical level compared to within fascicles of branches, and even further, could potentially account for the diverse clinical responses seen in patients undergoing VNS.

The dispersion of fibers seen in the IHC cross-sections of the cervical vagus nerve is unexpected. However, this observation was merely based on the recurrent laryngeal fascicles compared to the fascicles at cervical level and from those cross-sections. No tracing of fibers was performed, and there are still clusters of larger fibers visible at the cervical level. There may still be organization of organ-specific fibers at this level. This would be expected considering the pig vagus nerve was organized organotopically with respect to the three organs (Figure 15A), and human somatic nerves are organized somatotopically (1,35,36). On the other hand, the nodose ganglia house the cell bodies of the afferent fibers and the DVMN the efferent cell bodies. If the vagus nerve was reorganizing itself from organ-specific to fiber-specific, one would expect the clusters of larger fibers to be more pronounced at the cervical level and not dispersed. Tracing of fibers needs to be performed to provide these answers.

**Figure 15.**
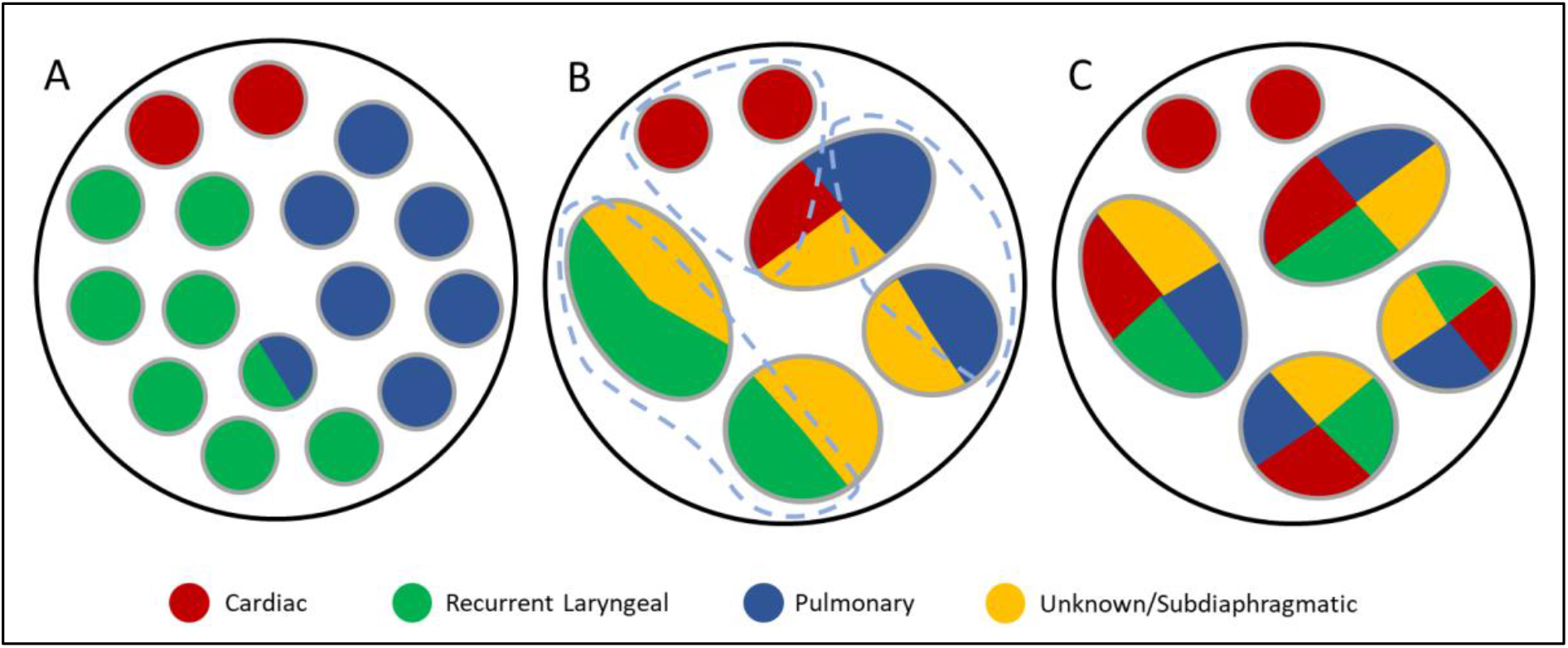
Didactic illustration of organization in the cervical vagus nerve of pigs and humans. A. Observed pig vagus nerve organization of cardiac, pulmonary and recurrent laryngeal fiber-containing fascicles (24). B. A potential organization of human cervical vagus nerve fascicles where there is spatial organization of fibers within the fascicles across the overall cross-section of the nerve, still allowing for selective activation of organ-specific fiber groups (encircled in grey dashed lines) despite distribution across multiple fascicles. C. A second potential organization of human cervical vagus nerve fascicles where there is no spatially separated localization of fibers across the total cross-section which would prove difficult to perform spatially-selective stimulation of a specific group of organ-specific fibers.

The fascicular anatomy of vagus nerve is highly complex and dynamic, and it appears even more so in the human vagus nerve. Mapping nerves using microCT is an effective technique to visualize and quantify fascicular organization. Understanding the nerve morphology and anatomy can assist in improving electrode cuff design as well as stimulation strategies. Given that humans have three to five times less fascicles, that are larger, than pig vagus nerves their fibers are expected to have higher activation thresholds (37,38). These analyses of fascicle dynamics, anatomy and organization within the human cervical vagus nerve provides further perspective into the morphology of the nerves and its implications on VNS efficacy. This data can contribute to computational modelling and *in vivo* studies for investigation of improved selectivity and vagal neuromodulation techniques. This will ultimately allow for improvement of the efficiency of treatment and the patients’ quality of life.

A clear limitation of this study was the fascicular resolution of the microCT scans which only allows for accurate and confident tracing of organ-specific fascicles prior to merger of other known organ-specific fascicles. This limited the tracing of organ-specific fascicles beyond clavicular level; however, microCT with segmentation still provided valuable information.

Specifically, microCT with iodine staining identifies fascicles from the rest of the soft tissue of the nerve; it does not enable visualisation of individual nerve fibres within the fascicles. In this work, I employed the following principle for assignment of function to rostral cervical fascicles. I identified the function of a distal peripheral branch by its proximity to the supplied end organ. I then traced the branch and respective fascicles rostrally. Providing the fascicles either did not diverge rostrally, only converged rostrally, or merged with the same fascicle type and subsequently diverged rostrally, it was logical to infer that the assigned organ function was unique. However, if two or more fascicles supplied by different organs merged and then diverged rostrally, it was unclear which organ function was subserved in the rostral diverged fascicles. It is possible that each fascicle was still organ specific, but it may also have been the case that nerve fibres had mixed so that each divergent rostral fascicle now was supplied by more than one organ. In either case, this organ supply cannot be identified from fascicle tracing alone. Based on the hypothesis that fibers pass between fascicles during this cross-over, when fascicles of two organ-specific types merged, the resulting merged fascicle was labelled as a mixed according to the organ-specific fascicles of origin. If that fascicle then continued proximally and diverged again, the resulting fascicles would maintain that mixed fascicle label. This erred on the side of caution as it may have been that some rostral divergent fascicles were actually still organ-specific. If all fascicles merge proceeding proximally, all resulting fascicles will be labelled as mixed fascicles, which is likely to be the case in human vagus nerves due to the presence of fewer fascicles.

Despite the limited (fascicular) resolution of microCT as mentioned above, by performing this microCT and tracing method, the following could be determined, quantified, analyzed, and subsequently compared between individuals: 1) the path of the fascicles within the nerve from entry into the vagus till cervical level, 2) the arrangement of fascicles at the cervical level, 3) the general pattern of fascicle movement, anastomoses and pathway in the nerve, 4) the number of fascicles for each branch and within the length of the vagus, 5) the number of and distances between branches to specific organs and effectors (general gross vagal anatomy), 6) the distances between first and last merging of fascicles of a specific organ-type within the vagus i.e., distances fascicles travel in a cable-like structure before merging with other fascicle types (fascicular morphology), and 7) the frequency of merging and splitting between fascicles of the same type and of different types i.e., the frequency of cross-over between fascicles, the plexiform structure of the nerves.

Other methods will be required to provide either fiber-specific tracing of the vagus nerve such as iDISCO (39), 3D-MUSE (40), or painstaking fiber tracing from serial IHC, or functional imaging of areas in the nerve with EIT or selective stimulation. Another limitation was not including the nodose ganglion in the dissections of the human vagus nerve. This would have allowed for tracing of fascicles bypassing the nodose, which would contain efferent fibers, and those originating from the nodose, which would contain afferent fibers. It may also have resulted in merger of fascicles and the same issue as above; however, this was indeed a short sight.

Performing histology and IHC at regular intervals allowed for morphological analyses and comparison with other studies. Additionally, this work served as protocol development for future human IHC studies and fiber analyses. Even by performing this at intervals as shown here, this will allow for calculation of the total number and ratio of fibers in the nerve, and in specific fascicles, and how this changes/shifts in proportion progressively at regular intervals in the nerve, with respect to efferent/afferent, myelinated/unmyelinated, and parasympathetic/sympathetic. In addition, it will be possible to determine the diameters of the fibers, and thereby, the numbers of various fiber types (A, B, C, etc.). If these methods were to be performed on sequential slices throughout the nerve the following could be determined: the path of the vagal fibers of the organ-specific fascicles from entry point in the vagus nerve to beyond the nodose ganglion to the brainstem. Specifically, the fibers could be traced throughout the merger and subsequently split of fascicles when progressing up the nerve (i.e. tracing organ-specific fibers, not fascicles, even when and after fascicles of various types merge and microCT cannot discern fascicle types, besides tracing a ‘merged’ fascicle for the rest of the nerve). In addition, a fiber map for all fascicles could be delineated at the cervical level and throughout the vagus nerve length, specific to organ and fiber type (efferent/afferent, presence of any hitchhiking sympathetic, etc.). Lastly, fiber tracing and microCT fascicle tracing a could be compared to determine any patterns that can be deduced/inferred from one to the other.

Choline Acetyltransferase (ChAT) is the enzyme that is responsible for biosynthesis of the neurotransmitter acetylcholine. The majority of acetylcholine is synthesized locally at nerve terminals where ChAT catalyzes the transfer of an acetyl group from acetyl coenzyme A to choline, a process that takes place in a single step. ChAT is expressed by cholinergic neurons in the central and peripheral nervous systems (41). Despite being a general parasympathetic histochemical stain – being classed as cholinergic-specific – it seems to have different sensitivities to motor (efferent) vs sensory (afferent) fibers; a study showed it is present predominantly in motor and scantly in sensory in a ratio of 8:1 (42). ChAT has been used frequently to differentiate between efferent and afferent fibers (42–44). With immunohistochemical markers such as this, and sometimes in a combination, the various neuron types can be identified. Specifically, motor markers include ChAT, cholinesterase, tyrosine kinase (TrkA), and agrin whilst sensory markers include calcitonin gene-related peptide (CGRP) and transient receptor potential vanillic acid subtype 1 (TRPV1), and annexin V (42,44–46). Sympathetic fibers can be distinguished from parasympathetic by using tyrosine hydroxylase (47). While many studies have used various combinations of IHC markers to distinguish fibers, such as motor vs sensory, no method seems to be standardized or largely adopted with new method development studies as recent as this year (2023). However, it seems probable to develop a combination of IHC stains to provide informative anatomical characterization of parasympathetic and, specifically, vagus nerves.

Future work will consist of fiber analysis of the cross-sections at regular intervals and from the branches including quantifying and measuring the fiber sizes, types, ratios, and their organization across the slice, and lastly, how this changes/shifts in proportion progressively at regular intervals in the nerve. It needs to be determined whether the fascicles at cervical level are still somewhat organized in a manner consistent with the pig vagus nerves (Figure 15A) and the human somatic nerves with fibers localized into a region of the whole vagus nerve cross-section, such as in Figure 15B, which would still allow for spatially selective stimulation of organ-specific fibers, or if there is little spatial organization of organ-specific fibers across the cross-section at cervical level at all as in Figure 15C. ChAT staining should be optimized for human nerves in the future. This would provide insight into the distribution of efferent fibers within the fascicles of the nerve. Additionally, double staining with NF and MBP should be done with fluorescence. This will allow for easier and more efficient visualization of the neuron versus the myelin sheath and will allow for quantification of numbers and sizes with readily available software that can detect fluorescence with ease. Tyrosine hydroxylase could also be detected for to see if any sympathetic fibers are present in the vagus nerve. Additionally, in order to determine functional organization in human vagus nerves, EIT and selective stimulation is being used in an ongoing *in vivo* clinical study. This will allow for localization of organ-specific functional activity within the cervical vagus nerve, and thereby determination of possible organization beyond structural imaging methods, and could ultimately allow for a patient-specific map to guide targeted stimulation for each individual.

## Conclusions

Similar to the somatic nervous system and the organotopic organization observed in pigs, it seems reasonable to postulate that despite organ-specific fascicles merging and anastomosing throughout proportions of the human vagus nerve, fibers within merged fascicles in the human vagus nerve at cervical level could still be somewhat arranged according to their supply to individual organs and possibly specific functions. Studies *in vivo* in human patients testing the electrophysiology of the nerve, stimulating spatially separated regions of the vagus and observing physiological readouts for different organ function, such as EIT and selective stimulation in (24) could assist with providing information beyond the limits of the fascicular resolution of microCT.

The superior cardiac fascicles remained separate at level of cuff placement which shows promise for selectivity of neuromodulation of the heart. This could benefit a number of conditions for which pharmaceuticals and other treatments are not sufficient in some cases, such as myocardial infarction, heart failure, atrial fibrillation (48–50).

Here, we have provided more morphological information of human vagus nerves to contribute to the minimal knowledge currently available and so contributing to the currently insufficient anatomical dataset that could assist with improvement of electrode cuff designs, stimulation strategies, and improvements to selective VNS and thereby accelerate the development of novel neuromodulation therapies for autonomic regulation.

## Abbreviations

ANS: Autonomic nervous system
CGRP: Calcitonin gene-related peptide
ChAT: Choline acetyltransferase
CNS: Central nervous system
CT: Computed tomography
d: Distinguishability
DI: Designate individual
EIT: Electrical impedance tomography
fps: Frames per second
GIT: Gastrointestinal tract
H&E: Haemotoxylin and Eosin
HTA: Human Tissue Authority
I: Iodine
MBP: Myelin basic protein
MicroCT: Micro-computed tomography
NBF: Neutral buffered formalin
NF: Neurofilament
NIH: National Institutes of Health
PNS: Peripheral nervous system
PRV: Pseudorabies virus
PsNS: Parasympathetic nervous system
ROI: Region of interest
sVNS: Selective vagus nerve stimulation
TH: Tyrosine hydroxylase
VNS: Vagus nerve stimulation

## Acknowledgements

The authors thank the Cancer Institute Pathology Lab, UCL, particularly Ayse Akarca and Teresa Marafioti, for the use of their labs and assistance with the automated staining machines, and lastly, who without, the human study would not have been possible, the Evelyn Cambridge Surgical Training Centre, A Cambridge University Health Partners Facility, with special thanks to HTA officer Dr Christopher Constant, for the loan of human tissue.

## Authors’ contributions

NT conceived and designed the study, performed dissections, collected and analyzed data, interpreted results and wrote the manuscript. SM performed dissections and edited the manuscript. KA and DH supervised the work and edited the manuscript. FI and PS provided resources for microCT imaging. All authors read and approved the final version of the paper.

## Availability of data and materials

The datasets used and/or analyzed during the current study are available from the corresponding author on request.

## Declarations

### Consent for publication

Not Applicable.

### Author details

^1^Department of Medical Physics and Biomedical Engineering, University College London; Gower Street, London WC1E 6BT, United Kingdom

^2^Electrochemical Innovations Lab, Department of Chemical Engineering, University College London; Gower Street, London WC1E 6BT, United Kingdom

## References

1. Bäumer P, Weiler M, Bendszus M, Pham M. Somatotopic fascicular organization of the human sciatic nerve demonstrated by MR neurography. Neurology. 2015 Apr 28;84(17):1782–7.

2. Shearer BM. The Morphology and Evolution of the Primate Brachial Plexus. 2019.

3. Thompson N, Mastitskaya S, Holder D. Avoiding off-target effects in electrical stimulation of the cervical vagus nerve: Neuroanatomical tracing techniques to study fascicular anatomy of the vagus nerve. Journal of Neuroscience Methods. 2019 Jun 26;325:108325.

4. Pelot NA, Goldhagen GB, Cariello JE, Musselman ED, Clissold KA, Ezzell JA, et al. Quantified Morphology of the Cervical and Subdiaphragmatic Vagus Nerves of Human, Pig, and Rat. Frontiers in Neuroscience. 2020;14:1148.

5. Rea P. Chapter 10 - Vagus Nerve. In: Rea P, editor. Clinical Anatomy of the Cranial Nerves [Internet]. San Diego: Academic Press; 2014 [cited 2018 Sep 19]. p. 105–16. Available from: http://www.sciencedirect.com/science/article/pii/B9780128008980000105

6. Ravagli E, Mastitskaya S, Thompson N, Iacoviello F, Shearing PR, Perkins J, et al. Imaging fascicular organization of rat sciatic nerves with fast neural electrical impedance tomography. Nature Communications. 2020 Dec 7;11(1):6241.

7. Plachta DTT, Gierthmuehlen M, Cota O, Espinosa N, Boeser F, Herrera TC, et al. Blood pressure control with selective vagal nerve stimulation and minimal side effects. Journal of Neural Engineering. 2014 May;11(3):036011.

8. Ardell JL, Rajendran PS, Nier HA, KenKnight BH, Armour JA. Central-peripheral neural network interactions evoked by vagus nerve stimulation: functional consequences on control of cardiac function. American Journal of Physiology-Heart and Circulatory Physiology. 2015 Sep 14;309(10):H1740–52.

9. Bai L, Mesgarzadeh S, Ramesh KS, Huey EL, Liu Y, Gray LA, et al. Genetic Identification of Vagal Sensory Neurons That Control Feeding. Cell. 2019 Nov 14;179(5):1129–1143.e23.

10. Rajendran PS, Nakamura K, Ajijola OA, Vaseghi M, Armour JA, Ardell JL, et al. Myocardial infarction induces structural and functional remodelling of the intrinsic cardiac nervous system. The Journal of Physiology. 2016;594(2):321–41.

11. Capllonch-Juan M, Sepulveda F. Modelling the effects of ephaptic coupling on selectivity and response patterns during artificial stimulation of peripheral nerves. PLOS Computational Biology. 2020 Jun 1;16(6):e1007826.

12. Sheheitli H, Jirsa VK. A mathematical model of ephaptic interactions in neuronal fiber pathways: Could there be more than transmission along the tracts? Netw Neurosci. 2020 Jul 1;4(3):595–610.

13. Isabella AJ, Stonick JA, Dubrulle J, Moens CB. Intrinsic positional memory guides target-specific axon regeneration in the zebrafish vagus nerve. Development. 2021 Sep 14;148(18):dev199706.

14. Rajendran PS, Challis RC, Fowlkes CC, Hanna P, Tompkins JD, Jordan MC, et al. Identification of peripheral neural circuits that regulate heart rate using optogenetic and viral vector strategies. Nat Commun [Internet]. 2019 Apr 26 [cited 2019 Jun 3];10. Available from: https://www.ncbi.nlm.nih.gov/pmc/articles/PMC6486614/

15. Mastitskaya S, Thompson N, Holder D. Selective Vagus Nerve Stimulation as a Therapeutic Approach for the Treatment of ARDS: A Rationale for Neuro-Immunomodulation in COVID-19 Disease. Front Neurosci [Internet]. 2021 [cited 2021 May 14];15. Available from: https://www.frontiersin.org/articles/10.3389/fnins.2021.667036/full

16. Mulders DM, de Vos CC, Vosman I, van Putten MJAM. The effect of vagus nerve stimulation on cardiorespiratory parameters during rest and exercise. Seizure. 2015 Dec 1;33:24–8.

17. Fitchett A, Mastitskaya S, Aristovich K. Selective Neuromodulation of the Vagus Nerve. Frontiers in Neuroscience. 2021 May;15:600.

18. Bonaz B, Bazin T, Pellissier S. The Vagus Nerve at the Interface of the Microbiota-Gut-Brain Axis. Front Neurosci [Internet]. 2018 Feb 7 [cited 2019 Oct 15];12. Available from: https://www.ncbi.nlm.nih.gov/pmc/articles/PMC5808284/

19. Breit S, Kupferberg A, Rogler G, Hasler G. Vagus Nerve as Modulator of the Brain–Gut Axis in Psychiatric and Inflammatory Disorders. Front Psychiatry [Internet]. 2018 Mar 13 [cited 2019 Aug 20];9. Available from: https://www.ncbi.nlm.nih.gov/pmc/articles/PMC5859128/

20. Browning KN, Verheijden S, Boeckxstaens GE. The Vagus Nerve in Appetite Regulation, Mood, and Intestinal Inflammation. Gastroenterology. 2017 Mar 1;152(4):730–44.

21. Koopman FA, Chavan SS, Miljko S, Grazio S, Sokolovic S, Schuurman PR, et al. Vagus nerve stimulation inhibits cytokine production and attenuates disease severity in rheumatoid arthritis. Proceedings of the National Academy of Sciences. 2016 Jul 19;113(29):8284–9.

22. Hammer N, Löffler S, Cakmak YO, Ondruschka B, Planitzer U, Schultz M, et al. Cervical vagus nerve morphometry and vascularity in the context of nerve stimulation - A cadaveric study. Scientific reports. 2018 May 22;8(1):7997.

23. Verlinden TJM, Rijkers K, Hoogland G, Herrler A. Morphology of the human cervical vagus nerve: Implications for vagus nerve stimulation treatment. Acta Neurologica Scandinavica. 2016 Mar 1;133(3):173–82.

24. Thompson N, Ravagli E, Mastitskaya S, Iacoviello F, Stathopoulou TR, Perkins J, et al. Organotopic organization of the porcine mid-cervical vagus nerve. Frontiers in Neuroscience [Internet]. 2023 [cited 2023 May 9];17. Available from: https://www.frontiersin.org/articles/10.3389/fnins.2023.963503

25. Schindelin J, Arganda-Carreras I, Frise E, Kaynig V, Longair M, Pietzsch T, et al. Fiji: an open-source platform for biological-image analysis. Nature Methods. 2012 Jul;9(7):676– 82.

26. O’Connor WN, Valle S. A combination Verhoeff’s elastic and Masson’s trichrome stain for routine histology. Stain Technol. 1982 Jul;57(4):207–10.

27. Yuan A, Rao MV, Veeranna null, Nixon RA. Neurofilaments at a glance. J Cell Sci. 2012 Jul 15;125(Pt 14):3257–63.

28. Deber CM, Reynolds SJ. Central nervous system myelin: structure, function, and pathology. Clin Biochem. 1991 Apr;24(2):113–34.

29. Sheehan DC, Hrapchak BB. Theory and practice of histotechnology. Columbus, Ohio: Battelle Press; 1987.

30. Hammer N, Löffler S, Cakmak YO, Ondruschka B, Planitzer U, Schultz M, et al. Cervical vagus nerve morphometry and vascularity in the context of nerve stimulation - A cadaveric study. Sci Rep [Internet]. 2018 May 22 [cited 2018 Oct 8];8. Available from: https://www.ncbi.nlm.nih.gov/pmc/articles/PMC5964190/

31. Hammer N, Glätzner J, Feja C, Kühne C, Meixensberger J, Planitzer U, et al. Human Vagus Nerve Branching in the Cervical Region. PLOS ONE. 2015 Feb 13;10(2):e0118006.

32. Stakenborg N, Gomez-Pinilla PJ, Verlinden TJM, Wolthuis AM, D’Hoore A, Farré R, et al. Comparison between the cervical and abdominal vagus nerves in mice, pigs, and humans. Neurogastroenterology & Motility. 2020;32(9):e13889.

33. Verlinden TJM, Rijkers K, Hoogland G, Herrler A. Morphology of the human cervical vagus nerve: implications for vagus nerve stimulation treatment. Acta Neurologica Scandinavica. 2016 Mar;133(3):173–82.

34. Upadhye AR, Kolluru C, Druschel L, Al Lababidi L, Ahmad SS, Menendez DM, et al. Fascicles split or merge every 560 microns within the human cervical vagus nerve. J Neural Eng [Internet]. 2022 [cited 2022 Oct 5]; Available from: http://iopscience.iop.org/article/10.1088/1741-2552/ac9643

35. Stewart JD. Peripheral nerve fascicles: Anatomy and clinical relevance. Muscle & Nerve. 2003;28(5):525–41.

36. Zill SN, Underwood MA, Rowley JC, Moran DT. A somatotopic organization of groups of afferents in insect peripheral nerves. Brain Research. 1980 Oct 6;198(2):253–69.

37. Pelot NA, Behrend CE, Grill WM. Modeling the response of small myelinated axons in a compound nerve to kilohertz frequency signals. J Neural Eng. 2017 Aug;14(4):046022.

38. Grinberg Y, Schiefer MA, Tyler DJ, Gustafson KJ. Fascicular Perineurium Thickness, Size, and Position Affect Model Predictions of Neural Excitation. IEEE Trans Neural Syst Rehabil Eng. 2008 Dec;16(6):572–81.

39. Renier N, Wu Z, Simon DJ, Yang J, Ariel P, Tessier-Lavigne M. iDISCO: A Simple, Rapid Method to Immunolabel Large Tissue Samples for Volume Imaging. Cell. 2014 Nov 6;159(4):896–910.

40. Kolluru C, Subramaniam A, Liu Y, Upadhye A, Khela M, Druschel L, et al. 3D imaging of the vagus nerve fascicular anatomy with cryo-imaging and UV excitation. In: Three-Dimensional and Multidimensional Microscopy: Image Acquisition and Processing XXVIII [Internet]. International Society for Optics and Photonics; 2021 [cited 2021 Apr 20]. p. 1164910. Available from: https://www.spiedigitallibrary.org/conference-proceedings-of-spie/11649/1164910/3D-imaging-of-the-vagus-nerve-fascicular-anatomy-with-cryo/10.1117/12.2577037.short

41. Oda Y. Choline acetyltransferase: the structure, distribution and pathologic changes in the central nervous system. Pathol Int. 1999 Nov;49(11):921–37.

42. Agarwal P, Bajaj J, Sharma D. Techniques for Differentiating Motor and Sensory Fascicles of a Peripheral Nerve—A Review. Indian Journal of Neurotrauma. 2020 Jun;17(01):28–32.

43. Settell ML, Pelot NA, Knudsen BE, Dingle AM, McConico AL, Nicolai EN, et al. Functional vagotopy in the cervical vagus nerve of the domestic pig: implications for the study of vagus nerve stimulation. J Neural Eng. 2020 Apr;17(2):026022.

44. Zhou X, Du J, Qing L, Mee T, Xu X, Wang Z, et al. Identification of sensory and motor nerve fascicles by immunofluorescence staining after peripheral nerve injury. Journal of Translational Medicine. 2021 May 13;19(1):207.

45. Riley DA, Sanger JR, Matloub HS, Yousif NJ, Bain JLW, Moore GH. Identifying motor and sensory myelinated axons in rabbit peripheral nerves by histochemical staining for carbonic anhydrase and cholinesterase activities. Brain Research. 1988 Jun 21;453(1):79–88.

46. Meng X, Lu L, Wang H, Liu B. Differentiation between the motor and sensory fascicles of the peripheral nerves from adult rats using annexin V-CdTe-conjugated polymer. Neurology India. 2011 Jan 5;59(3):333.

47. Weihe E, Depboylu C, Schütz B, Schäfer MKH, Eiden LE. Three Types of Tyrosine Hydroxylase-Positive CNS Neurons Distinguished by Dopa Decarboxylase and VMAT2 Co-Expression. Cell Mol Neurobiol. 2006;26(0):659–78.

48. De Ferrari GM, Schwartz PJ. Vagus nerve stimulation: from pre-clinical to clinical application: challenges and future directions. Heart Fail Rev. 2011 Mar 1;16(2):195–203.

49. Sabbah HN, Ilsar I, Zaretsky A, Rastogi S, Wang M, Gupta RC. Vagus Nerve Stimulation in Experimental Heart Failure. Heart Fail Rev. 2011 Mar;16(2):171–8.

50. Yao Y, Kothare MV. Model Predictive Control of Selective Vagal Nerve Stimulation for Regulating Cardiovascular System. In: 2020 American Control Conference (ACC). 2020. p. 563–8.

