## Supplementary Information for "Human vagus nerve fascicular anatomy: a microCT segmentation and histological study"

**IHC Protocol:**

1. *Immerse in TBS buffer (Sigma 94158) or BOND Wash Solution (Leica Biosystems AR9590) – here on out referred to as TBS buffer*
2. *Immerse in hydrogen peroxide in TBS for 15 minutes*
3. *Wash slides with TBS with 3 changes of 2 minutes each*
4. *Immerse in 5% donkey serum (Sigma D9663) for 30 minutes*
5. *Rinse with water*
6. *Add diluted primary antibody(ies) for 1 hour (NF and MBP together)*
    a. *For NF: anti-neurofilament heavy polypeptide antibody ab8135 1:1000*
    b. *For MBP: anti-myelin basic protein antibody ab7349 1:4000*
7. *Wash slides with TBS with 3 changes of 2 minutes each*
8. *Add secondary biotin labelled antibody diluted in TBS 1:1000 for 1 hour (donkey anti-rabbit IgG H&L (Biotin) ab6801)*
9. *Wash slides with TBS with 3 changes of 2 minutes each*
10. *Add tertiary streptavidin/biotin complex in TBS for 30 min*
11. *Wash slides with TBS with 3 changes of 2 minutes each*
12. *Add DAB solution (Sigma D3939) for 5 minutes*
13. *Wash with distilled water*

14. *Add secondary HRP labelled antibody diluted in TBS 1:1000 for 1 hour (donkey anti-rat IgG H&L (HRP) ab102182)*
15. *Wash slides with TBS with 3 changes of 10 minutes each*
16. *Add Vector NovaRed Substrate Kit (Vector Laboratories SK-4800) tertiary solution in TBS for 30 min*
17. *Wash slides with TBS with 3 changes of 10 minutes each*
18. *Add the Vector NovaRed substrate for 5 minutes*
19. *Wash with distilled water*
20. *Add Harris's hematoxylin (diluted 1:1) for 1 minute*
21. *Wash with water for 3 minutes*
22. *Dehydrate through alcohols into xylene*
23. *Remove from the IHC machine and place in water*
24. *Coverslip slides with DPX*

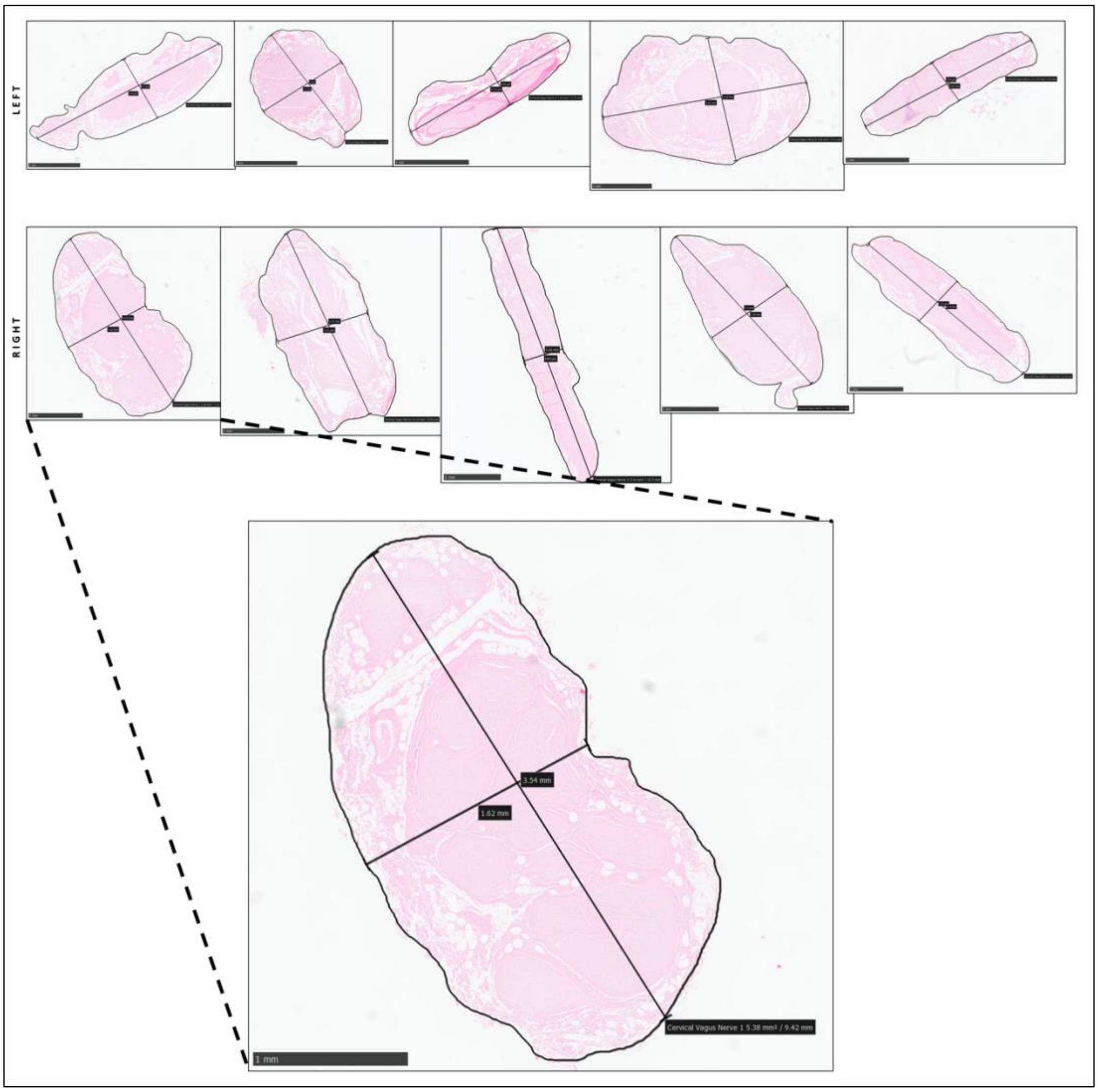


**Supplementary Figure 1. Histology cross section measurements.** Cross sections from mid-cervical level of five left and five right human vagus nerves (n=10) stained with H&E displaying two diameter (short and long) and circumference measurements for each for morphological analysis.

### Supplementary Table 1. Morphological analysis of the human cervical vagus nerve (n=10)

| *Human Nerve* | *Side* | *Circumference (mm)* | *Area (mm2)* | *Short Diameter (mm)* | *Long Diameter (mm)* | *Mean Diameter* | *No. of fascicles* | *No. of fascicle bundles* |
|---|---|---|---|---|---|---|---|---|
| 1 | R | 9.42 | 5.38 | 1.62 | 3.52 | 2.57 | 20 | 11 |
| 2 | L | 9.92 | 3.82 | 1.3 | 3.94 | 2.62 | 14 | 5 |
| 3 | R | 8.59 | 4.26 | 1.51 | 3.19 | 2.35 | 7 | 4 |
| 4 | L | 5.89 | 2.4 | 1.5 | 2 | 1.75 | 6 | 4 |
| 5 | R | 10.5 | 3.16 | 0.699 | 4.66 | 2.6795 | 12 | 4 |
| 6 | L | 5.75 | 1.43 | 0.544 | 2.45 | 1.497 | 9 | 3 |
| 7 | R | 9.37 | 4.44 | 1.59 | 3.5 | 2.545 | 11 | 7 |
| 8 | L | 9.41 | 5.82 | 2.09 | 3.43 | 2.76 | 11 | 6 |
| 9 | R | 9.16 | 3.62 | 1.03 | 3.74 | 2.385 | 8 | 8 |
| 10 | L | 6.8 | 1.93 | 0.73 | 2.93 | 1.83 | 8 | 7 |
| ***Mean*** | | 8.48 | 3.63 | 1.26 | 3.34 | 2.30 | 10.60 | 5.90 |
| ***Std Dev*** | | 1.70 | 1.43 | 0.50 | 0.75 | 0.44 | 4.12 | 2.42 |
| ***Mean L*** | | 7.55 | 3.08 | 1.23 | 2.95 | 2.09 | 9.60 | 5.00 |
| ***Std Dev L*** | | 1.98 | 1.77 | 0.62 | 0.77 | 0.56 | 3.05 | 1.58 |
| ***Mean R*** | | 9.41 | 4.17 | 1.29 | 3.72 | 2.51 | 11.60 | 6.80 |
| ***Std Dev R*** | | 0.69 | 0.85 | 0.41 | 0.56 | 0.14 | 5.13 | 2.95 |

#### Supplementary Table 2. Fascicle and fascicle bundle counts from regular intervals of the vagus nerve trunk and organ-specific branches

| *Row 1 – Fascicles* | *1cm* | *3cm* | *5cm* | *9cm* | *13cm* | *17cm* | *21cm* | ***Mean*** | ***Std Dev*** |
|---|---|---|---|---|---|---|---|---|---|
| *Row 2 – Bundles* | | | | | | | | | |
| *Row 3 – Fascicles* | *Superior cardiac* | *Recurrent laryngeal* | *Inferior cardiac* | *Pulmonary* | | | | | |
| *Row 4 – Bundles* | | | | | | | | | |
| Nerve 1 | 11 | 20 | 20 | 18 | 24 | 15 | N/A | 18.00 | 4.52 |
| | 9 | 11 | 17 | 12 | 12 | 10 | N/A | 11.83 | 2.79 |
| | 4 | / | 3 | 3 | | | | | |
| | 3 | / | 3 | 3 | | | | | |
| Nerve 2 | 8 | 14 | 14 | 22 | 12 | N/A | N/A | 14.00 | 5.10 |
| | 6 | 5 | 10 | 15 | 11 | N/A | N/A | 9.40 | 4.04 |
| | 2 | 7 | 2 | / | | | | | |
| | 2 | 6 | 2 | / | | | | | |
| Nerve 3 | 7 | 6 | 8 | 9 | 21 | 13 | N/A | 10.67 | 5.61 |
| | 4 | 5 | 4 | 8 | 12 | 8 | N/A | 6.83 | 3.13 |
| | 3 | / | 2 | 5 | | | | | |
| | 3 | / | 2 | 5 | | | | | |
| Nerve 4 | 6 | 4 | 7 | 31 | 7 | 7 | N/A | 10.33 | 10.19 |
| | 4 | 2 | 3 | 14 | 6 | 6 | N/A | 5.83 | 4.31 |
| | 6 | 8 | 4 | 6 | | | | | |
| | 6 | 8 | 4 | 4 | | | | | |
| Nerve 5 | 15 | 12 | 11 | 12 | 17 | 6 | 8 | 11.57 | 3.78 |
| | 7 | 4 | 9 | 11 | 14 | 4 | 8 | 8.14 | 3.97 |
| | 2 | 10 | 1 | 7 | | | | | |
| | 2 | 10 | 1 | 7 | | | | | |
| Nerve 6 | 9 | 11 | 9 | 16 | 1 | 7 | N/A | 8.83 | 4.92 |
| | 3 | 8 | 3 | 13 | 1 | 6 | N/A | 5.67 | 4.37 |
| | 1 | 8 | 2 | 6 | | | | | |
| | 1 | 7 | 2 | 6 | | | | | |
| Nerve 7 | 11 | 11 | 20 | 9 | 16 | 14 | N/A | 13.50 | 4.04 |
| | 7 | 7 | 16 | 7 | 13 | 8 | N/A | 9.67 | 3.88 |
| | 4 | 5 | 3 | 4 | | | | | |
| | 4 | 4 | 3 | 3 | | | | | |
| Nerve 8 | 3 | 4 | 11 | 23 | 14 | 7 | N/A | 10.33 | 7.47 |
| | 2 | 4 | 6 | 22 | 12 | 4 | N/A | 8.33 | 7.53 |
| | 4 | 7 | 1 | / | | | | | |
| | 4 | 6 | 1 | / | | | | | |
| Nerve 9 | 8 | 9 | 12 | 11 | 28 | 8 | 25 | 14.43 | 8.42 |

| | | | | | | | | | |
|---|---|---|---|---|---|---|---|---|---|
| | 5 | 7 | 11 | 9 | 25 | 7 | 19 | 11.86 | 7.31 |
| | 2 | 4 | 3 | 12 | | | | | |
| | 2 | 4 | 3 | 8 | | | | | |
| Nerve 10 | 5 | 6 | 8 | 11 | 9 | 1 | 8 | 6.86 | 3.24 |
| | 5 | 5 | 7 | 7 | 7 | 1 | 5 | 5.29 | 2.34 |
| | 3 | 9 | / | / | | | | | |
| | 3 | 4 | / | / | | | | | |
| ***Mean*** | 8.30 | 9.70 | 12.00 | 16.20 | 14.90 | 8.67 | 13.67 | 11.85 | 5.73 |
| | 5.20 | 5.80 | 8.60 | 11.80 | 11.30 | 6.00 | 10.67 | 8.29 | 4.37 |
| | 3.10 | 7.25 | 2.33 | 6.14 | | | | | |
| | 3.00 | 6.13 | 2.33 | 5.14 | | | | | |
| ***Std Dev*** | 3.43 | 5.01 | 4.71 | 7.28 | 8.14 | 4.50 | 9.81 | 3.20 | 2.25 |
| | 2.10 | 2.53 | 5.02 | 4.59 | 6.25 | 2.69 | 7.37 | 2.42 | 1.74 |
| | 1.45 | 1.98 | 1.00 | 2.91 | | | | | |
| | 1.41 | 2.17 | 1.00 | 1.95 | | | | | |

**Supplementary Table 3. Trichrome cross sections at regular intervals along the trunk of the vagus and from organ-specific branches**

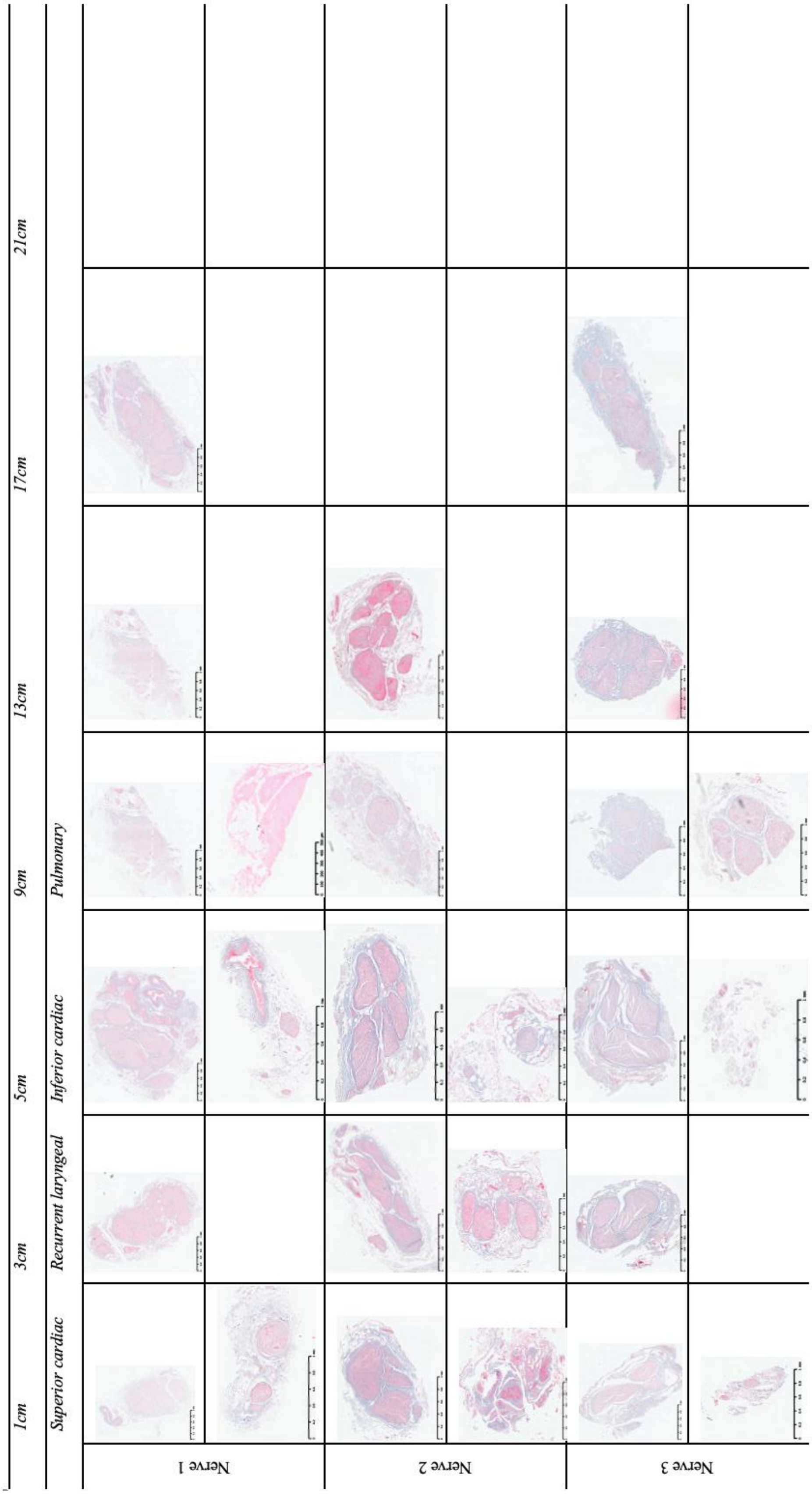

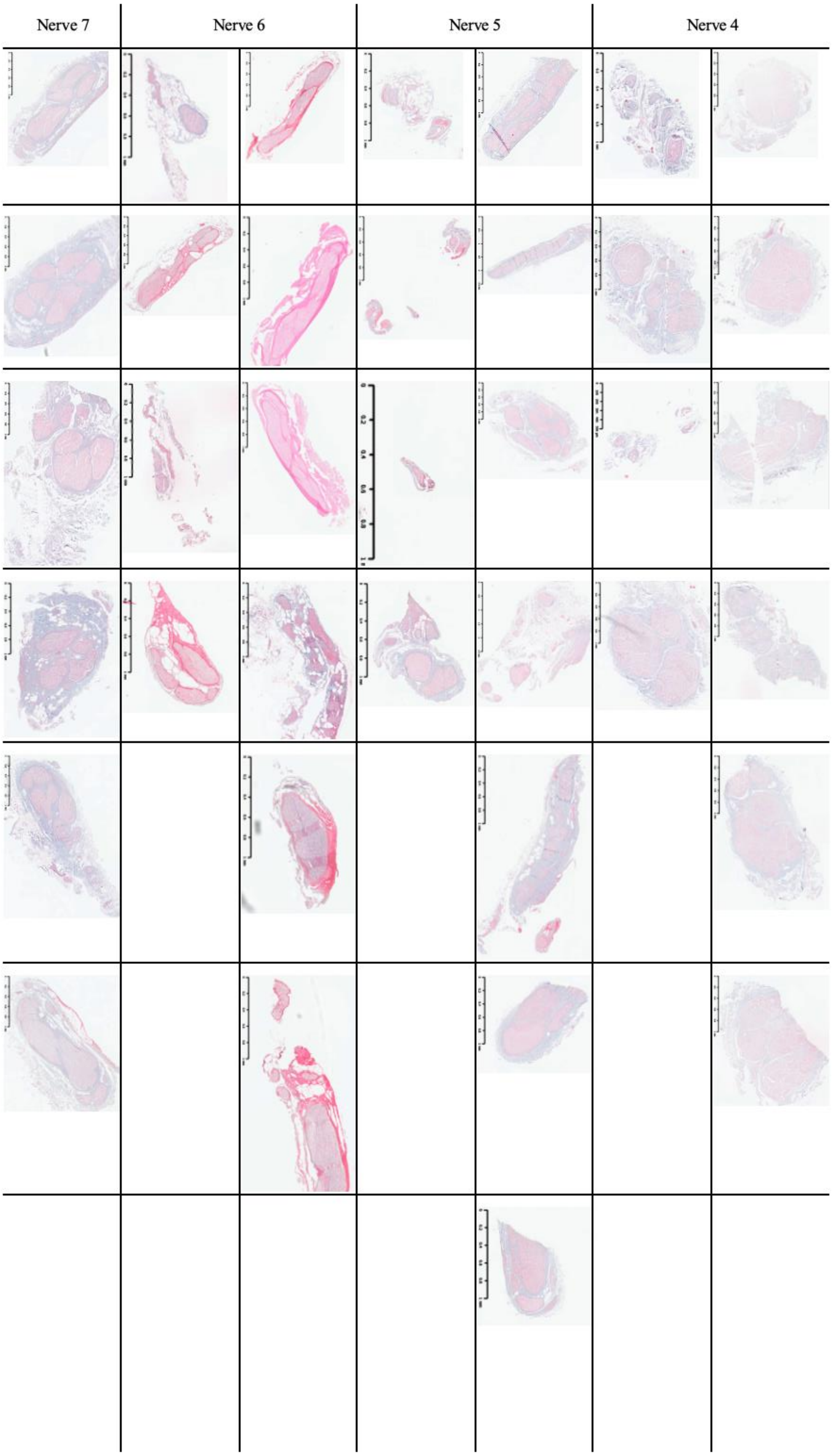
Nerve 7
Nerve 6
Nerve 5
Nerve 4

| Nerve 10 | Nerve 9 | Nerve 8 |
| --- | --- | --- |

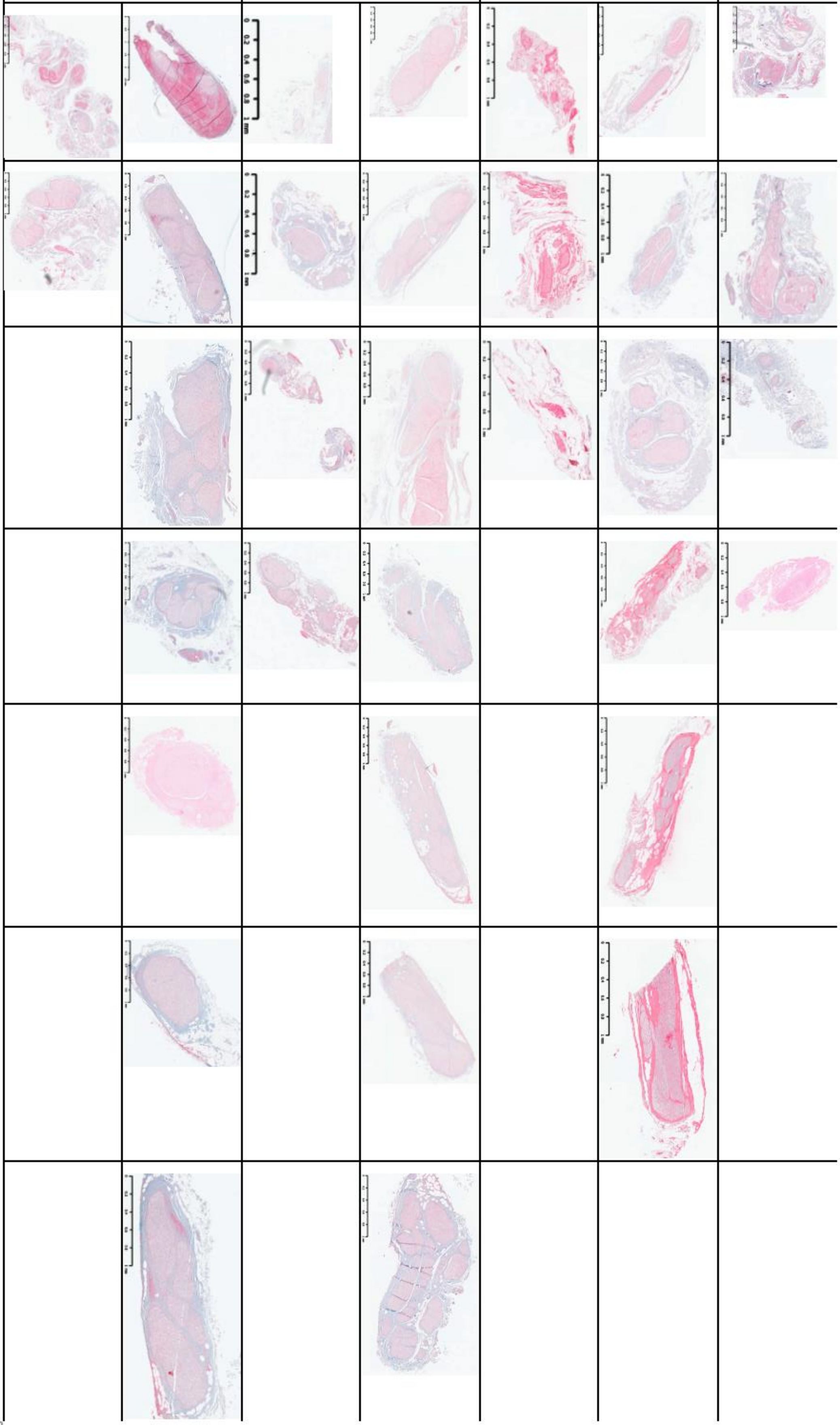

**Supplementary Table 4. Human vagus nerve branch measurements: distances from cervical level to organ-specific branches and distance between branches exiting the vagus nerve**

| *Distance from cervical level (cm)* | | | | | | | | | | |
|---|---|---|---|---|---|---|---|---|---|---|
| *Nerve* | 1 | 2 | 3 | 4 | 5 | 6 | 7 | 8 | 9 | 10 |
| *Side* | R | L | R | L | R | L | R | L | R | L |
| *Superior cardiac (SC)* | 4.5 | 3.75 | 8 | 3 | 7 | 7 | 10.5 | 8 | 8 | 4 |
| *Recurrent laryngeal (RL)* | 10.5 | 15 | 10 | 15 | 14 | 13.5 | 13.5 | 15.5 | 13.5 | 17.5 |
| *Inferior cardiac 1 (IC1)* | 10.5 | 14.5 | 10 | 15 | 14 | 13.5 | 13.5 | 15 | 13.5 | 17 |
| *Inferior cardiac 2 (IC2)* | 11 | 15 | 10.5 | 15.5 | 15 | 13.5 | 14 | 15.5 | 14 | 17.5 |
| *Pulmonary 1 (P1)* | 18 | 17 | 16.5 | 16 | 20 | 16 | 19 | 18 | 20.5 | 19 |
| *Pulmonary 2 (P2)* | 18.5 | 17.5 | 17 | 18 | 21 | 18 | 21 | 19 | 21 | 21 |
| *Pulmonary 3 (P3)* | 19 | N/A | 17.5 | N/A | 22 | 19 | N/A | N/A | 21.5 | N/A |

| | ***Mean*** | ***Std Dev*** | ***Mean R*** | ***Std Dev R*** | ***Mean L*** | ***Std Dev L*** |
|---|---|---|---|---|---|---|
| *SC* | 6.38 | 2.43 | 7.60 | 2.16 | 5.15 | 2.21 |
| *RL* | 13.80 | 2.24 | 12.30 | 1.89 | 15.30 | 1.44 |
| *IC1* | 13.65 | 2.08 | 12.30 | 1.89 | 15.00 | 1.27 |
| *IC2* | 14.15 | 2.11 | 12.90 | 2.01 | 15.40 | 1.43 |
| *P1* | 18.00 | 1.62 | 18.80 | 1.60 | 17.20 | 1.30 |
| *P2* | 19.20 | 1.64 | 19.70 | 1.86 | 18.70 | 1.40 |
| *P3* | 19.80 | 1.89 | 20.00 | 2.12 | 19.00 | N/A |

| *Distance between branches (cm)* | | | | | | | | | | |
|---|---|---|---|---|---|---|---|---|---|---|
| *Nerve* | 1 | 2 | 3 | 4 | 5 | 6 | 7 | 8 | 9 | 10 |
| *Side* | R | L | R | L | R | L | R | L | R | L |
| *SC - RL* | 6 | 11.25 | 2 | 12 | 7 | 6.5 | 3 | 7.5 | 5.5 | 13.5 |
| *RL - IC1* | 0 | -0.5 | 0 | 0 | 0 | 0 | 0 | -0.5 | 0 | -0.5 |
| *IC1 - IC2* | 0.5 | 0.5 | 0.5 | 0.5 | 1 | 0 | 0.5 | 0.5 | 0.5 | 0.5 |
| *RL - P1* | 7.5 | 2 | 6.5 | 1 | 6 | 2.5 | 5.5 | 2.5 | 7 | 1.5 |
| *IC2 - P1* | 7 | 2 | 6 | 0.5 | 5 | 2.5 | 5 | 2.5 | 6.5 | 1.5 |
| *P1 - P2* | 0.5 | 0.5 | 0.5 | 2 | 1 | 2 | 2 | 1 | 0.5 | 2 |
| *P2 - P3* | 0.5 | N/A | 0.5 | N/A | 1 | 1 | N/A | N/A | 0.5 | N/A |

| | ***Mean*** | ***Std Dev*** | ***Mean R*** | ***Std Dev R*** | ***Mean L*** | ***Std Dev L*** |
|---|---|---|---|---|---|---|
| *SC - RL* | 7.43 | 3.78 | 4.70 | 2.11 | 10.15 | 3.01 |
| *RL - IC1* | -0.15 | 0.24 | 0.00 | 0.00 | -0.30 | 0.27 |
| *IC1 - IC2* | 0.50 | 0.24 | 0.60 | 0.22 | 0.40 | 0.22 |
| *RL - P1* | 4.20 | 2.52 | 6.50 | 0.79 | 1.90 | 0.65 |

| | | | | | | |
|---|---|---|---|---|---|---|
| *IC2 - P1* | 3.85 | 2.31 | 5.90 | 0.89 | 1.80 | 0.84 |
| *P1 - P2* | 1.20 | 0.71 | 0.90 | 0.65 | 1.50 | 0.71 |
| *P2 - P3* | 0.70 | 0.27 | 0.63 | 0.25 | 1.00 | N/A |

**Supplementary Video 1. Example of anastomoses (splitting and merging) of fascicles in the human vagus nerve**

https://youtube.com/shorts/0GgmLg2mN-0?feature=share
